# Biophysical determinants of mutational robustness in a viral molecular fitness landscape

**DOI:** 10.1101/172197

**Authors:** Manasi A. Pethe, Aliza B. Rubenstein, Dmitri Zorine, Sagar D. Khare

**Author notes:** Rutgers, The State University of New Jersey, Piscataway NJ 08854.

## Abstract

Biophysical interactions between proteins and peptides are key determinants of genotype-fitness landscapes, but an understanding of how molecular structure and residue-level energetics at protein-peptide interfaces shape functional landscapes remains elusive. Combining information from yeast-based library screening, next-generation sequencing and structure-based modeling, we report comprehensive sequence-energetics-function mapping of the specificity landscape of the Hepatitis C Virus (HCV) NS3/4A protease, whose function – site-specific cleavages of the viral polyprotein – is a key determinant of viral fitness. We elucidate the cleavability of 3.2 million substrate variants by the HCV protease and find extensive clustering of cleavable and uncleavable motifs in sequence space indicating mutational robustness, and thereby providing a plausible molecular mechanism to buffer the effects of low replicative fidelity of this RNA virus. Specificity landscapes of known drug-resistant mutations in the protease are similarly clustered, indicating mutational robustness in both the enzyme and its substrates. Our results highlight the key and constraining role of molecular-level energetics, acting as a “biophysical capacitor”, in shaping plateau-like fitness landscapes predicted by viral quasispecies theory.

## Introduction

RNA viruses, e.g., influenza, Hepatitis C virus (HCV) and Human Immunodeficiency virus (HIV), are under a heavy mutational load due to the extremely high error-rates of their RNA polymerases(Domingo and Holland, 1997; Holland et al., 1982; Lauring et al., 2013). As a result of this low replication fidelity, these viruses exist as a population of variants called quasispecies(Andino and Domingo, 2015; Eigen, 1993), even within a single host individual (Cristina et al., 2007). While this genetic diversity and a large population size is believed to increase viral adaptive potential against antiviral therapies(Elde et al., 2012; Goldberg et al., 2012; Wilke et al., 2001), low replication fidelity may also lead to too many mutations, causing an “error catastrophe” and extinction(Eigen, 2002; Lauring and Andino, 2010). The underlying biomolecular structures and interactions in the virus must, therefore, be robust to genetic variability such that they provide a buffer against the deleterious impacts of a high mutational load (Elena et al., 2006; Masel and Siegal, 2009). Tawfik and co-workers have hypothesized that viral proteins possess “gradient robustness” in which individual mutations have small and largely additive effects on stability leading to a slower loss of function compared to “threshold robustness” exhibited by proteins in general (Tokuriki et al., 2009). It has been argued that mutational robustness may itself promote adaptiveness if the number of phenotypes accessible to a variant through mutation is smaller than the total number of phenotypes possible(Draghi et al., 2010; Wilke and Adami, 2003). How is mutational robustness encoded at the molecular level in RNA viruses such as HCV? How is structural integrity and interaction fidelity maintained in the face of a large mutational load, and what, if any, are the limits imposed by the underlying molecular interactions on mutational robustness and adaptive potential? The degeneracy of the genetic code, the thermodynamic and kinetic stabilities of RNA and proteins, and the presence of molecular chaperones, may all contribute to the robustness of the *structures* of individual viral biomolecules (Lauring et al., 2013). However, how viral protein-based *interactions*, especially those that are critical for viral propagation, encode “fuzziness” (Tokuriki et al., 2009) leading to mutational robustness at the molecular level is not well understood.

At the molecular level, the balance between mutational robustness and functional plasticity is encapsulated in the notion of molecular fitness landscapes(Smith, 1970), which are high-dimensional maps that relate the function of individual biomolecular variants to their functional and/or evolutionary fitness (de Visser and Krug, 2014; Wright, 1931). Analysis of mutational trajectories on these landscapes provides insight into the constraints placed on evolution by the physiochemical properties of biomolecules, allowing, in principle, reconstruction as well as forward prediction of molecular evolution (Bridgham et al., 2006; Harms and Thornton, 2013; Kondrashov and Kondrashov, 2015; Romero and Arnold, 2009; Weinreich et al., 2006). The molecular fitness landscape has long been theoretically postulated (Smith, 1970) and recent empirically determined sequence-function mappings of proteins (Bandaru et al., 2017; Firnberg et al., 2014; Fowler et al., 2010; Hietpas et al., 2011; Kim et al., 2013; McLaughlin et al., 2012; Podgornaia and Laub, 2015; Sarkisyan et al., 2016; Wrenbeck et al., 2017) have enabled the partial construction of fitness landscapes. These reconstructed landscapes permit testing of possible evolutionary scenarios and provide insight into properties such as mutational robustness and non-additivity (epistasis) of mutational effects (Blanquart and Bataillon, 2016; Breen et al., 2012; Harms and Thornton, 2013; Hartl, 2014; Sailer and Harms, 2017a; Thyagarajan and Bloom, 2014; Weinreich et al., 2013; Wu et al., 2016). Empirical sequence-function relationships also enable biomolecular engineering for new or improved functions (Jenson et al., 2017; Klesmith et al., 2015; McLaughlin et al., 2012; Tinberg et al., 2013; Whitehead et al., 2012).

Typically, sequence-function mapping of proteins and protein-protein interactions described above involves partial enumeration of the possible sequence diversity (for example, all single mutations and a subset of double mutations at a large number of protein residue positions) and high-throughput functional evaluation coupled with deep sequencing(Fowler and Fields, 2014; Klesmith et al., 2017; Reich et al., 2015). Statistical and/or biophysical models can be used to make inferences about the regions of sequence space not sampled(Jenson et al., 2017; Klesmith et al., 2017). However, comprehensive construction of the fitness landscape requires enumeration and evaluation of the complete sequence diversity (all higher-order mutations at all residue positions). Laub and coworkers have pioneered studies in which the entire combinatorial diversity is experimentally sampled, albeit at a smaller number of positions(Aakre et al., 2015; Podgornaia and Laub, 2015). The astronomical size of sequence space, however, makes the comprehensive experimental evaluation of sequence-function landscapes with any one experimental approach difficult. Computational biophysical methods may, in principle, assist in creation and analysis of functional and fitness landscapes (Rodrigues et al., 2016). Indeed, evolutionary landscapes of simple protein models, such as lattice models, have been extensively investigated using biophysical evolutionary theory and computational simulations(Bloom et al., 2004; Bornberg-Bauer and Chan, 1999; DePristo et al., 2005; Ding and Dokholyan, 2006; Drummond and Wilke, 2008; Echave and Wilke, 2017; Manhart and Morozov, 2015; Sailer and Harms, 2017b; Sikosek and Chan, 2014; van Nimwegen et al., 1999; Yang et al., 2012), and deep connections with population genetics theories have been discovered (Bershtein et al., 2017; Echave and Wilke, 2017; Serohijos and Shakhnovich, 2014). While pioneering and crucial insights have been obtained in these studies, chemically realistic atomic-resolution structure-based elucidation of functional landscapes has not been performed so far, due both to high computational cost as well as inaccuracies in simulation force fields which preclude accurate biophysical evaluation of mutational effects on protein interactions.

Here, we use a combination of experimental (biochemical) and computational techniques to elucidate the specificity landscape of the interaction between HCV NS3/4A protease enzymes and its substrates. This enzyme-substrate interaction is key for viral maturation as it cleaves exclusively at four specific sites in the viral polyprotein (Fig. 1A) to release individual non-structural proteins(Scheel and Rice, 2013), and also mediates inactivation of key human immunity proteins(Meylan et al., 2005). The cleavage specificity of the protease is thus a key determinant of viral fitness, and its proper functioning includes negative specificity – the lack of cleavage of non-canonical sites on the viral protein and of most host cell proteins (Fig. 1A). The molecular interactions underlying both positive and negative specificities must be robust to mutations as the HCV virus RNA polymerase has a high error-rate (Powdrill et al., 2011), but how and whether this robustness is encoded in the protease-substrate interactions is not known. Using yeast surface display, next-generation sequencing and a machine-learning approach which combines features from experimental data and atomistic computational simulations (utilizing the Rosetta and Amber force fields) that we recently developed (Pethe et al., 2017; Rubenstein et al., 2017), we construct the specificity landscape (with cleavability assignments made for 3.2 million substrate pentapeptide sequences) of the HCV NS3/4A protease and three of its known drug-resistant variants(Romano et al., 2012). We demonstrate that energetic features of protease-substrate interactions inherently encode mutational robustness, and that the connectivity patterns in the specificity landscape may act as a “biophysical capacitor” for maintaining protease function in the face of high mutational load.

**Figure 1:**
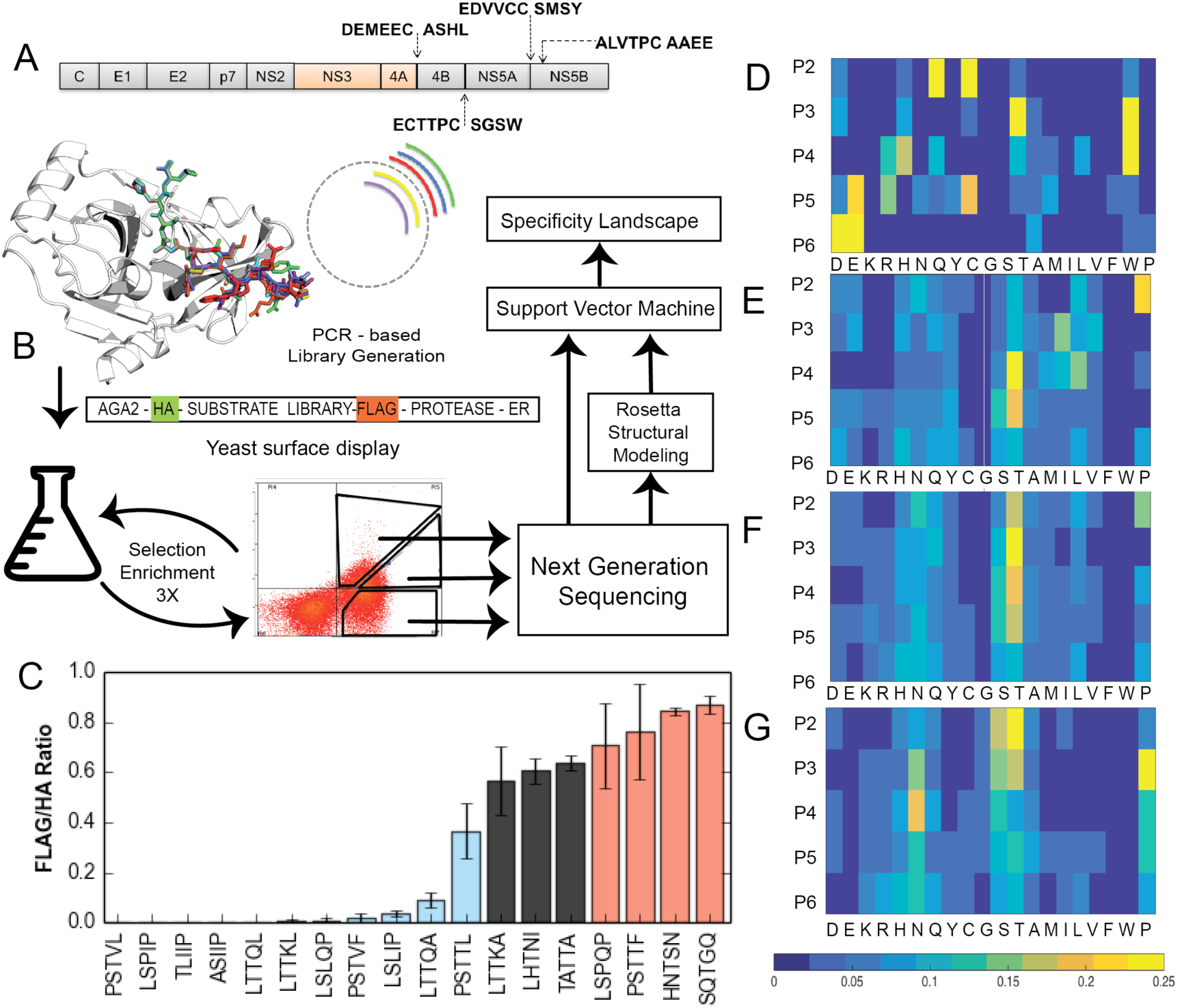
(A)The HCV viral polyprotein depicting marked biological cleavage sites for the HCV NS3 protease (B) Overview of the experimental and computational workflow. (C) Validation of FACS gates for cleaved, partially cleaved, and uncleaved sequences using the yeast surface display assay (D) Sequences taken from *in vivo* samples of FICV patients (8726) as compared to (E) sequences determined by our assay as cleaved (7472), (F) sequences determined by our assay as partially cleaved (8737), and (G) sequences determined by our assay as uncleaved (14702).

## Results

HCV NS3/4A protease is known to cleave four canonical cleavage sites on the hepatitis C viral polyprotein (Fig. 1A), causing a cascade of viral assembly and maturation events (Lindenbach and Rice, 2013). These cleavages (and a lack of cleavage of other parts of the polyprotein) are thus, critical for viral fitness. The high mutational load on the HCV polyprotein can lead to sequence variation in both the protease and substrate regions (Geller et al., 2016a). At the protein level, the distribution of mutational effects in a folded protein (protease) are modulated by both the thermodynamic stability and function (binding and cleavage), while the substrate regions, which are found in flexible linker regions of the HCV polyprotein (with no known tertiary structure) and connect component proteins, do not have a native tertiary structure. Therefore, we reasoned that a more direct sequence-cleavability mapping can be made for diversity in the substrate region without the need to additionally deconvolute the contribution from stability effects on tertiary structure. Secondly, it is more feasible to enumerate and evaluate by sequencing the substrate combinatorial diversity due to its shorter length (~7 residues) compared to the protease. Therefore, we mapped the viral protease-substrate interaction landscape for the HCV NS3/4A protease by considering all possible pentapeptide sequence combinations in its sequence recognition site at positions P6 through P2 following the Schechter and Berger nomenclature(Schechter and Berger, 1967). Positions P1 and P1’, between which the scissile bond is present, were maintained as C and A, respectively, in this study. In the rest of this paper, we refer to individual pentapeptide patterns (e.g., the canonical cleavage sites DEMEE, EDVVC, ECTTP, ALVTP) and omit the identity of the P1,P1’ residues.

### Exploration of the (P6-P2) specificity landscape of the HCV NS3/4A protease reveals a diverse specificity profile

To mimic the viral intrachain arrangement of substrate libraries and the protease, we utilized a modified version of the assay described by Iverson, Georgiou and co-workers(Yi et al., 2013) as depicted in Figure 1B. A mutagenic library was created incorporating degenerate codons at P6-P2 specificity defining substrate positions (Benatuil et al., 2010; Kowalsky et al., 2015). In our assay, substrates are transported to the surface of yeast cells in a cleavage-dependent manner: the degree of cleavage after induction of the fusion protein is estimated by measuring the relative levels of substrate-flanking FLAG and HA tags using fluorescent, labeled antibodies. We have previously used this assay to test known and novel substrates of the HCV protease (Pethe et al., 2017). A first round of yeast surface display assay and Fluorescence Assisted Cell Sorting (FACS) was performed with an inactive protease variant (S139A) to select for high expression of library variants, for removing sequences containing stop codons in the substrate region, and to deplete substrate sequences that are cleaved by yeast ER proteases (Li et al., 2017).

The resulting substrate variants from the pre-selection were subjected to rounds of yeast surface display assay and Fluorescence Assisted Cell Sorting (FACS) with an active protease containing construct to select cleaved, partially cleaved and uncleaved variants using three sorting gates (Fig. 1B) based on the relative levels of anti-HA and anti-FLAG fluorescence values (FLAG/HA ratio, ranging between 0, for completely cleaved, and 1, for completely uncleaved). Sorting gates were defined based on the distribution of populations observed for known cleaved and uncleaved sequences (Pethe et al., 2017). This procedure was coupled with rounds of growth and selection to improve signal:noise for variants in each pool. Sequence profiles of the isolated functional variants were determined using next-generation sequencing technology (Illumina NextSeq). Analysis of unique sequences in all sequenced pools showed that we identified a total of ~1.3 million sequences corresponding to ~30% of the possible amino acid diversity (3.2 million; Supplementary Information). Analysis of sequencing and biological duplicates as well as overlap between the sequence pools was used to determine a count threshold (raw count 11) to remove noise from the sequencing data (Methods and Supplementary Figure 1). Based on these criteria, we identified 7472, 8737 and 14702 unique pentapeptide sequences in the cleaved, partially cleaved and uncleaved pools. In parallel, we performed Rosetta simulations on all 3.2 million sequences in the P6-P2 region, and machine learning with Support Vector Machines to predict the complete protease-substrate interaction landscape using sequence information procured from the aforementioned library and Rosetta-generated energetic features (Figure 1B).

Several novel substrates identified from the three variant populations were tested as clonal populations in the yeast surface display assay system (Figure 1C, Supplementary Figure 2) to validate that individual sequences fall into the gates used for selection from the library (Supplementary Figure 3A-C). A subset of these sequences was also tested *in vitro* to ensure that the cleavage properties observed in the yeast system were reproduced with purified protease and substrates (Supplementary Figure 3D).

We next analyzed the profiles of substrates in each pool. For the cleaved sequence pool, the obtained substrate sequence ensemble has greater diversity compared to substrates identified from viral genomes sequenced from patient populations (Methods, Figure 1D). For example, we observe that a more diverse subset of amino acids is tolerated at substrate positions P6 and P5 in our cleaved and partially cleaved pools (Figure 1E, F) whereas the patient isolated genomes display a high enrichment of Asp and Glu specifically at these positions. Another notable difference observed was the enrichment of small hydrophilic residues (Figure 1E, F), Ser (at P5) and Thr (at P4) in the cleaved and partially cleaved populations, in contrast to enrichment at P3 and P2 in the uncleaved population (Figure 1G). Strikingly, we found prolines enriched at position P2 in the cleaved and partially cleaved populations and at P3 in the uncleaved populations, which corresponds well with the fact that 2 out of 4 canonical cleaved sequences have proline at P2 (ECTTP, ALVTP). While some of the above trends are also reflected in the sequences we tested in the course of our method validation (Fig. 1C), it is evident that individual positional enrichments cannot be directly used to predict the pool assignments of individual sequences. For example, His is enriched at P6 in the cleaved sequence pool, however the sequence HNTSN is experimentally determined to be in the uncleaved pool (Fig. 1C, Supplementary Figure 2). While individual positional preferences of amino acids are useful, these results clearly indicated that molecular recognition between the protease and substrate pools is highly (sequence) context-dependent. We concluded that interactions between amino acids at various substrate positions (mediated possibly through interaction networks in the protease) influence the cleavability, thereby motivating the need for an analysis of the determined specificity landscape using properties of whole pentapeptide sequences.

### Clustering among cleaved, partially cleaved and uncleaved substrates

To visualize the pattern of functionally labeled sequence space of the experimentally derived substrates, we generated a force-directed graph (Fig. 2A) (Amat, 2016; Jacomy et al., 2014) in which each node represents a sequence and is colored according to the functional pool to which it belongs. Nodes are connected by an edge if they differ by one amino acid. Cleaved substrates exhibit significant clustering in comparison to partially cleaved and uncleaved substrates, which are well distributed on the interaction landscape (Fig. 2A). To examine the landscape in greater detail around the cleaved sequences, we generated a sub-graph of the cleaved sequences (Fig. 2B). We identified four clusters in this graph using the Gephi (Amat, 2016) modularity algorithm and determined corresponding profiles for each cluster. One identified cluster is clearly related to a canonical substrate, DEMEE. The other three clusters appear to have similarities with the other three canonical substrates (ALVTP, ECTTP, and EDVVC) but are less distinct from each other compared to the DEMEE cluster. These results indicate that the four canonical cleaved sites in the viral polyprotein are all members of mutationally robust clusters. Single amino acid changes within the cluster lead to other cleaved sequences, thereby buffering the impact of the heavy mutational load on the virus.

**Figure 2:**
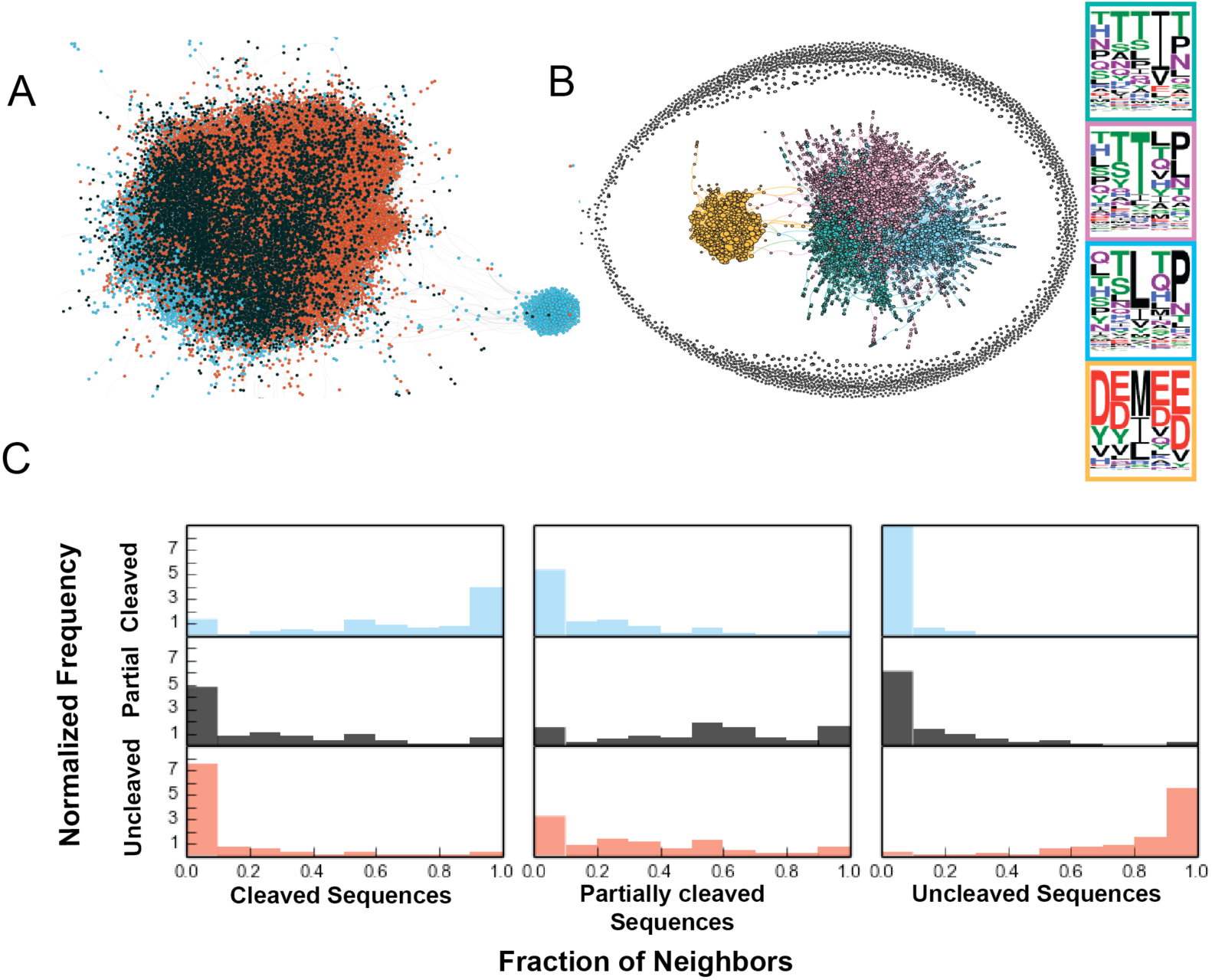
(A) Force-directed graph of amino acid sequence space. Blue nodes are cleaved, red are uncleaved, and black is partially cleaved. Edges connect nodes that are within one hamming distance of each other. (B) Force-directed graph of cleaved sequences. Colors denote clusters, which are shown as specificity profiles outlined in the same color as the corresponding cluster (C) Frequency of neighbors for cleaved, partially cleaved and uncleaved sequences denoting cleaved neighbors shown in blue bars uncleaved neighbors depicted in red and partially cleaved neighbors depicted in black bars.

To determine if this clustering behavior observed in the cleaved sequence pool is also found in the partially cleaved and uncleaved pools, we calculated the fraction of neighbors for sequences with neighbors that belong to the same functional pool. We find that similar to cleaved sequences, uncleaved sequences are also most frequently surrounded by uncleaved neighbors indicating clustering behavior for this functional pool as well. On average, cleaved sequence neighbors are 66.4% cleaved, and uncleaved sequence neighbors are 83.3% uncleaved. Partially cleaved sequences are the least clustered among the three pools, having on average 53% neighbors belonging to the same pool. These distributions indicate that in the specificity landscape, clusters of partially cleaved sequences surround clusters of cleaved and uncleaved ones.

To delineate how the three functional populations, which appear to be individually clustered in sequence space, are connected to each other through amino acid substitutions, we used the PageRank metric (Brin and Page, 1998). This metric predicts the likelihood of reaching a node given a random walk on the substrate specificity landscape starting from a chosen sequence. Strikingly, partially cleaved substrates have higher pageranks (Supplementary Figure 4A) than either cleaved or uncleaved substrates, indicating that they are most likely to be reached on long unbiased evolutionary trajectories starting from the canonical cleaved sequence DEMEE, the sequence that was used as the template for library generation. These connectivity patterns imply that partially cleaved node clusters may act as an evolutionary buffer on the substrate landscape, thereby enhancing robustness.

The graph generated by the experimentally derived sequences is incomplete (~30,000 nodes out of the 3.2 million possible). To test if the observed clustering and PageRank distributions are an artifact of the limited sampling in the experiment, we generated a control random graph (Supplementary Figure 5A) with the same number of nodes and edges, but having a randomly rewired connectivity. Both partially cleaved and uncleaved sequences are found to have higher pageranks than cleaved sequences in this random graph, indicating that the higher pageranks of partially cleaved sequences than cleaved and uncleaved sequences in the original experimental graph is significant.

### Energetic features derived from Rosetta modeling enable reconstruction of the complete protease-pentapeptide substrate landscape

While the experimentally-derived populations of the cleaved, partially cleaved and uncleaved sequences revealed striking clustering patterns in sequence space, it is not clear if these connectivity patterns would be preserved in a complete graph containing the complete diversity at five positions (3.2 million sequences). Therefore, to predict cleavability of all possible 3.2 million sequences in the interaction landscape, we used a Support Vector Machine (SVM)-based method that we developed previously (Pethe et al., 2017). Briefly, each sequence was threaded onto a bound complex based on a modeled near-attack conformation a crystal structure of the protease, and the complex was then relaxed to maintain favorable catalytic geometry. Energy evaluation of each of the 3.2 million complexes was performed using Rosetta and Amber simulation packages. A binary classification (cleaved/uncleaved) SVM was trained on a subset of experimentally identified sequences that passed a more stringent threshold of enrichment compared to the unselected pool in our assay (1817 cleaved and 3605 uncleaved sequences) as well as sequences identified by Shiryaev *et al* (Shiryaev et al., 2012) for a total of 7342 unique sequences. Training features consisted of structure-based features (energies of interaction) and sequence-based features (see Methods, Supplementary Figure 6A). We initially cross-validated the SVM on the training set using an 80:20 split with 100 iterations, which yielded an average AUROC of 0.96 (Supplementary Figure 6B) indicating high recapitulation of training data (a perfect performance would lead to an AUROC of 1). We then used the SVM to predict cleaved and uncleaved labels for the remaining 3,192,658 sequences. These predictions have a precision of 0.96 at a recall level of 0.91 for an overall accuracy of 0.96 (Supplementary Figure 6B) for the experimentally-derived assignments that were left out of the training set (~10000 sequences). We experimentally validated cleavage predictions for six substrates as clonal populations using the yeast assay and find good agreement with the SVM-based predictions (Fig. 3A). We visualized a sub-graph (force directed) of predicted cleaved sequences, present at a distance > 2 from the hyper-plane constructed by the SVM (Supplementary Figure 6C). The experimentally identified cleaved sequences are recapitulated well, and distributed evenly across the predicted cleaved population.

**Figure 3:**
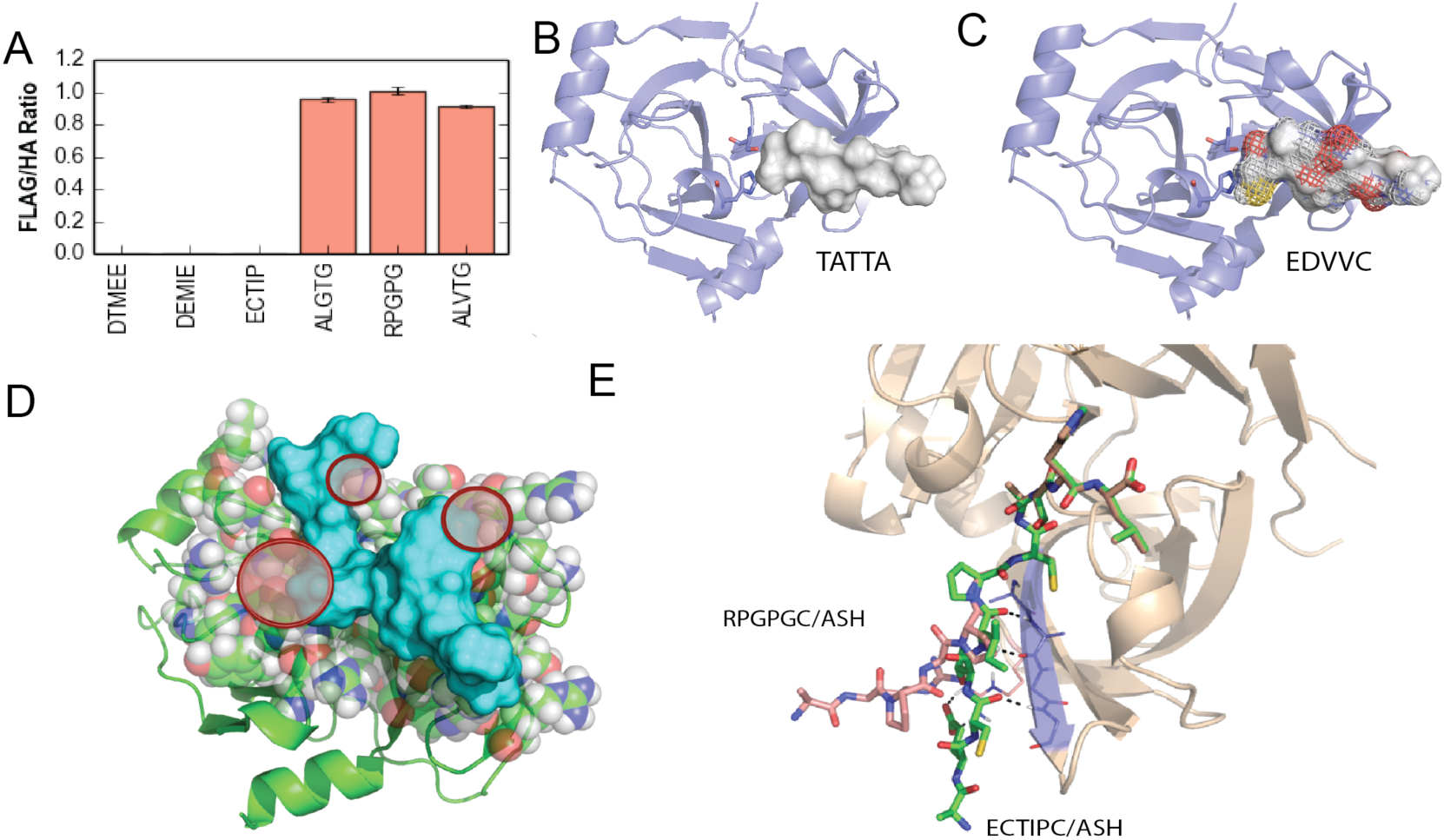
(A) Validation assay performed for three predicted cleaved and uncleaved sequences using a yeast surface display based technique.(B) and (C) depict the volume occupied by TATTA and EDVVC, EDVVC occupies an optimal volume, making good contacts with the protease residue side chains. TATTA fits in the available space but does not make optimal contacts, thus resulting in suboptimal interaction energetics making TATTA a suboptimal substrate. (D) Peptide (surface shown in blue) “FWWPM” sterically clashing against the protease chain. (E) Structure of two models, ECTIP (cleaved) and RPGPG (uncleaved).

### Structural and energetic bases for observed specificity patterns

Having obtained and validated predictions of cleavability by combining experimental and computational data, we turned structural models of protease-substrate complexes to obtain insight into the underlying structural basis of observed specificity patterns. For example, a comparative analysis of the partially cleaved substrate ‘TATTA’ and canonical substrate ‘EDVVC’ reveals that the former, composed of small residues does not completely occupy the substrate cavity volume (Figure 3B, C) whereas ‘EDVVC’ occupies the entire cavity. The lack of voids at the interface and several hydrogen bonds formed by the canonical lead to better binding (Binding interaction energy = -80.2 Rosetta energy units (Reu), as opposed to -77.5 Reu for TATTA), resulting in better cleavage for this substrate. Similarly, models of the uncleaved sequence FWPPM (Figure 3D) reveals that the side chains are found to have steric clashes with the protease side chains. Apart from sidechain-based interaction patterns, models also capture backbone conformational changes that affect the orientation of the substrate in the active site. For example, in the model corresponding to the sequence RPGPG(uncleaved), the proline present at P3 in RPGPG (Fig. 3E) bends the peptide chain away from the protease, resulting in breaking of the crucial backbone hydrogen bond patterns that are characteristic of protease-substrate interactions (Tyndall et al., 2005).

Structural analysis also allows rationalization of non-additive (epistatic) patterns between amino acid substitutions. We detected the presence of both positive and negative epistasis in our experimental data, and further investigated two cases (Figure 4A). We examined a predicted negative epistasis pathway (Figure 4B), where single-mutant P at position P4 and single-mutant Q at position P3 both result in a cleaved substrate but the double-mutant PQ at position P3-P4 is uncleaved. We measured the mutual information (Figure 4C; Methods) between positions P3 and P4 in the experimentally derived cleaved sequence pool and found that both L at P3 and Q at P4 (corresponding to sequence LSLQP) and P at P3 and I at P4 (corresponding to sequence LSPIP) are correlated, indicating that these two amino acid preferences are found in the experimentally-derived cleaved population at a higher incidence than expected by their individual incidence.

**Figure 4:**
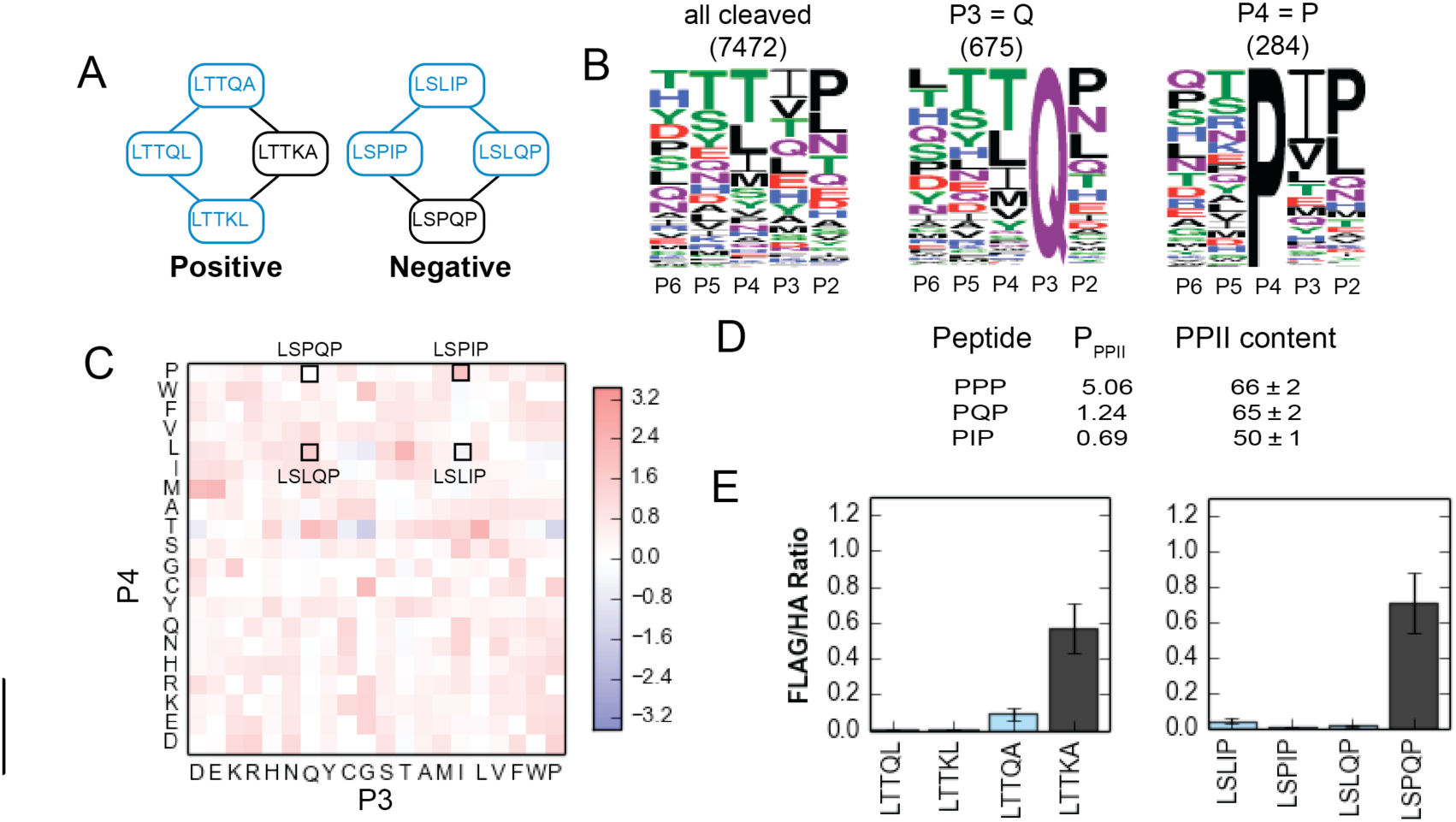
Structural basis underlying epistasis found on the interaction landscape.(A) Examples of positive and negative epistasis. Cleaved sequences are highlighted in blue, partially cleaved in red. (B) Specificity profiles for entire cleaved set (left), sequences with glutamine at P3 (middle), and proline at P4 (right). (C) Heatmap of correlations between positions 3 and 4, as measured by mutual information. (D) Polyproline II structure propensity of peptides (see text). (E) Experimental validation of the sequences in both positive and negative epistatic pathways, performed using yeast surface display. Blue bars indicate sequences that are expected to be cleaved and black bars indicate sequences that are expected to be partially cleaved.

However, the correlation for P at P4 and Q at P3 (corresponding to sequence LSPQP) is low, suggesting that the PQ pattern is depleted in the cleaved sequence population. Structurally, the sequence LSPQP (Figure 4D) may have an increased PPII (polyproline-II) helix propensity (Kelly et al., 2001; Vila et al., 2004), causing the substrate to twist out of a catalysis-competent binding conformation in our models. The PPII helix propensity for the sequence LSPIP is lower thus resulting in retention of the extended substrate binding conformation that is favorable for catalysis(Tyndall et al., 2005). Thus, analysis of models of individual substrates provides atom-resolution insights into how the underlying biophysics of molecular recognition by the protease shapes the observed specificity landscapes, including non-additive effects.

Having validated (Figure 4E) these examples of double-mutant epistatic networks, we investigated the double mutant epistatic networks present in the experimental data, and found that the majority of these epistatic networks (60.7%) involved cleaved and partially cleaved sequences only. The preponderance of epistatic networks at the cleaved/partially cleaved boundary indicates that the boundary between cleaved and partially cleaved sequences is more rugged than the boundary between cleaved and uncleaved sequences, further highlighting the role of partially cleaved sequences as a biophysical buffer in sequence space.

### Mutational robustness and possible evolutionary trajectories in the experimentally-determined and computationally reconstructed landscape

Having computed the entire P6-P2 specificity landscape, we next examined the connectivity patterns between cleaved and uncleaved sequences in this reconstructed landscape. As with the experimentally determined landscape, the reconstructed landscape also shows clear evidence of clustering between cleaved and uncleaved nodes (Supplementary Figure 4E-I), indicating that mutational robustness extends to regions of sequence space not covered in our library, and is an essential feature of this protease-substrate interface. As our SVM-based approach is a binary classification scheme, partially cleaved sequences are classified in either cleaved or uncleaved pools. Attempts to build a 3-way classifier failed due both to the noise from the experiments as well as difficulty in estimating small energy differences in Rosetta simulations. Further improvements in each methodology may allow the prediction of partially cleaved sequences.

As the Hepatitis C virus is subject to a considerable amount of evolutionary drift, we investigated the impact of the pathways of drifting on the landscape on maintaining function. For the experimentally determined landscape, we calculated the number of mutations from each canonical sequence to the functional boundary and plotted the fraction of cleaved substrates that can be reached at each step (Supplementary Figure 4E). The curves for both DEMEE and EDVVC reach a small initial plateau and then rise sharply, indicating that both are surrounded by a cluster of cleaved sequences and then must bridge a largely non-functional region of the graph to reach the rest of the cleaved sequences, whereas the curves for both ALVTP and ECTTP rise steadily, indicating that the topology surrounding these sequences is less rugged.

Both the reconstructed and experimentally-derived landscapes feature several novel “cleaved” sequence patterns (defined as >3 substitutions away from a canonical recognition motif). To investigate if these novel sequences can be reached, as an example, we generated a sub graph of the sequence space connecting the canonical cleaved sequences (DEMEE, EDVVC, ECTTP, ALVTP) with each other as well as the novel cleaved sequences, e.g., PSTVF (Fig. 5A). Analysis of all inter-node shortest paths on these networks shows that there exist many paths between canonical and novel sequences that do not include uncleaved nodes (viable paths) while some paths involve traversal of at least one predicted uncleaved node (unviable paths; Figure 5B). All canonical sequences are more highly connected to each other than to any of the novel sequence motifs, suggesting that the latter may be “kinetically” less accessible during evolutionary drifts. We calculated the fraction of non-viable paths between canonical sequences and compared it to the fraction of non-viable paths between canonical sequences and novel sequences. The latter shows a higher, albeit still small, fraction of non-viable paths (Fig. 5B). We also find that those novel cleaved sequences that have a higher fraction of cleaved neighbors (higher degree) are more likely to have a higher fraction of viable trajectories to canonical nodes. (Fig. 5C). Thus, it appears that the higher single mutational robustness of a given novel sequence is correlated with its ability to be reachable from/to canonical sequences that are at least three amino acid substitutions away in sequence space. Further contributions from codon usage in the host context may modulate the reachability of different substrates by making some amino acid changes even less likely. Our analysis above leaves out these contributions to selectively delineate the impact of amino acid-level effects.

**Figure 5:**
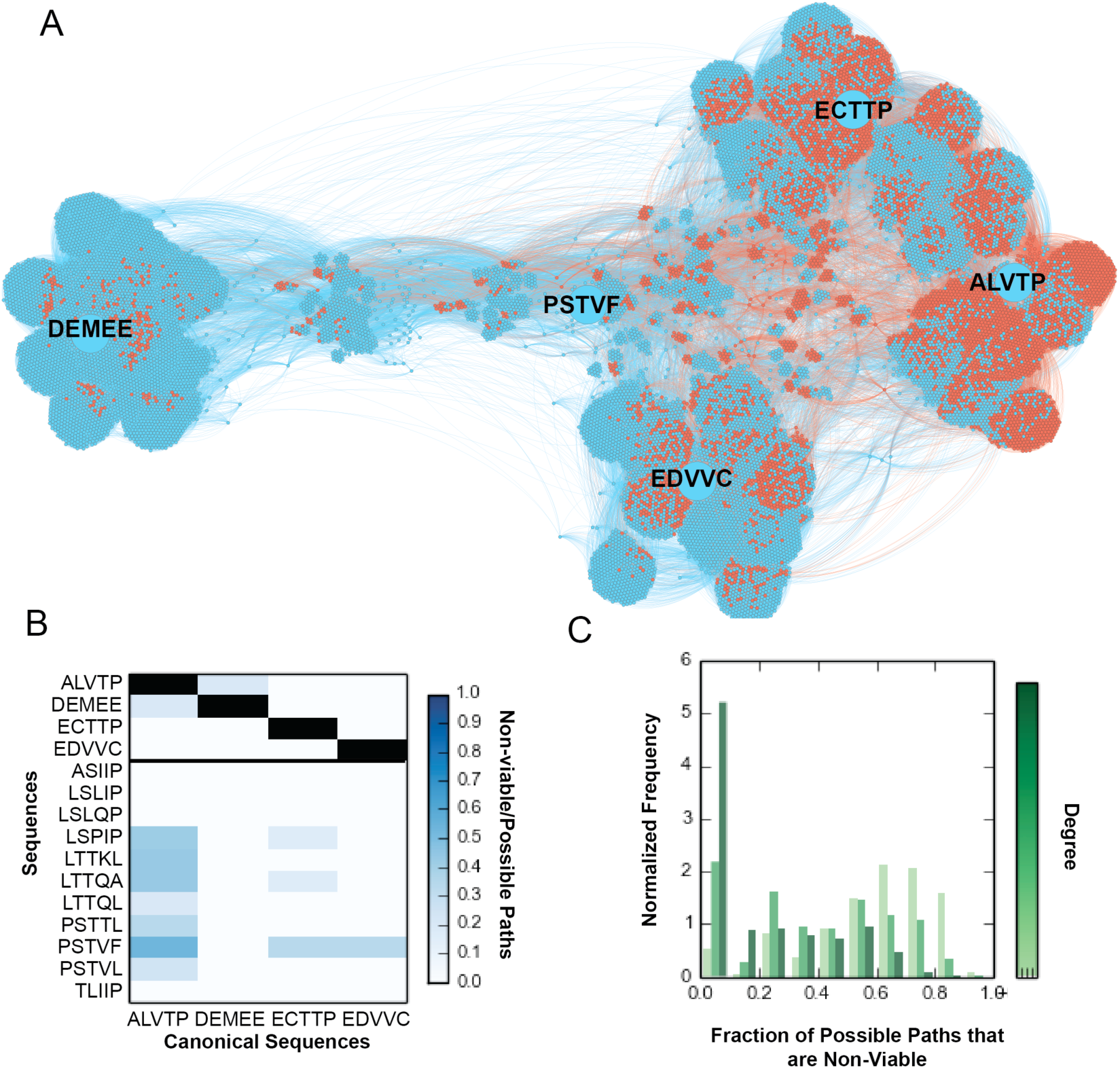
(A) A force directed interaction graph between the five canonical sequences - DEMEE, ECTTP, EDVVC, ALVTP and the novel cleaved sequence PSTVF(depicted by large blue nodes). The graph depicts neighbors of all intermediate sequences between PSTVF and all canonical sequences. The cleaved sequences in the interaction pathways are denoted by blue nodes and the uncleaved are denoted by red. (B) The fraction of uncleaved nodes present in the shortest paths from both canonical sequences and novel sequences to all canonical sequences. (C) Degree vs. fraction of the shortest paths uncleaved between all novel sequences and all canonical sequences.

### Protease specificity landscape may contribute to negative selection

Sequences of patient-derived genomes indicate that the HCV virus is under strong negative selection (Campo et al., 2008; Cuypers et al., 2015). Although the underlying mechanisms are not well understood, several factors have been invoked to explain the observation of a low dN/dS ratio (number of non-synonymous to synonymous substitutions in the genome) in the patient-derived populations including intrahost competition between quasispecies, and immune evasion(Skums et al., 2015). Given the centrality of the protease in viral maturation, we asked if maintenance of cleavability (and uncleavability) in different parts of the polyprotein also contributes to negative selection, and what, if any, are the limits imposed by the recognizability of different polyprotein regions by the protease on their variability.

As our reconstructed landscape provides information on all pentapeptide sequence combinations (followed by Cys-Ala), we asked if overlapping pentapeptides in the other parts of the polyprotein (apart from the known cleavage sites) are likely to be cleaved, especially if they acquired a Cys-Ala pattern in the two immediately downstream residues (thereby acquiring the necessary heptapeptide pattern that would be cleaved). If several regions of the polyprotein are poised to be cleaved upon acquisition of the Cys-Ala motif, an error catastrophe may ensue upon increasing the mutational load. We performed a genome-wide comparison of patient derived sequences with sequences predicted as cleaved by our SVM classifier. Each viral genome (Cuypers et al., 2015) was split into overlapping 5-mer peptide sequence fragments using a one-residue sliding window method. These 5-mers were compared to the pentapeptide sequences predicted by our approach as cleaved. If the patient-derived pentapeptide sequence was found in the cleaved pool, we calculated the minimum nucleotide mutational distance of the successive two residues from the DNA sequences that code for ‘CA’ and ‘CS’ which are known to be the canonical P1-P1’ sites favoring cleavage by the HCV NS3/4A protease (Fig. 1A). The results (Figure 6A,B, Supplementary Figure 7A-H) indicate that the majority (~70%) of patient-derived translated pentapeptides are are found the uncleaved pool. Of the remaining (~30%) 5-mer sequences that are identified as potentially cleavable (if they acquire a CA or CS as the following two amino acids), 74.1% pentapeptides from all genotypes of the virus require more than three nucleotide changes to acquire a ‘CA’ or ‘CS’ at the P1-P1’ sites (Supplementary Figure 7A-H). The avoidance of acquisition of a cleavable sequence in other regions of the protein, made feasible by codon usage, may thus, contribute to the previously described negative selection pressure on the HCV genome(Campo et al., 2008), and may be reflected in the measured low dN/dS rates in the non-structural regions of the protein(Cuypers et al., 2015). Additional avoidance of non-productive cleavage may also result from altered dynamics (Fuchs et al., 2014) and/or the post-translational structural context of the potentially cleavable regions – these may be buried (inaccessible to the protease) or adopt secondary structures that are incompatible with the extended conformation required to fit in the protease active site(Barkan et al., 2010; Julien et al., 2016).

**Figure 6:**
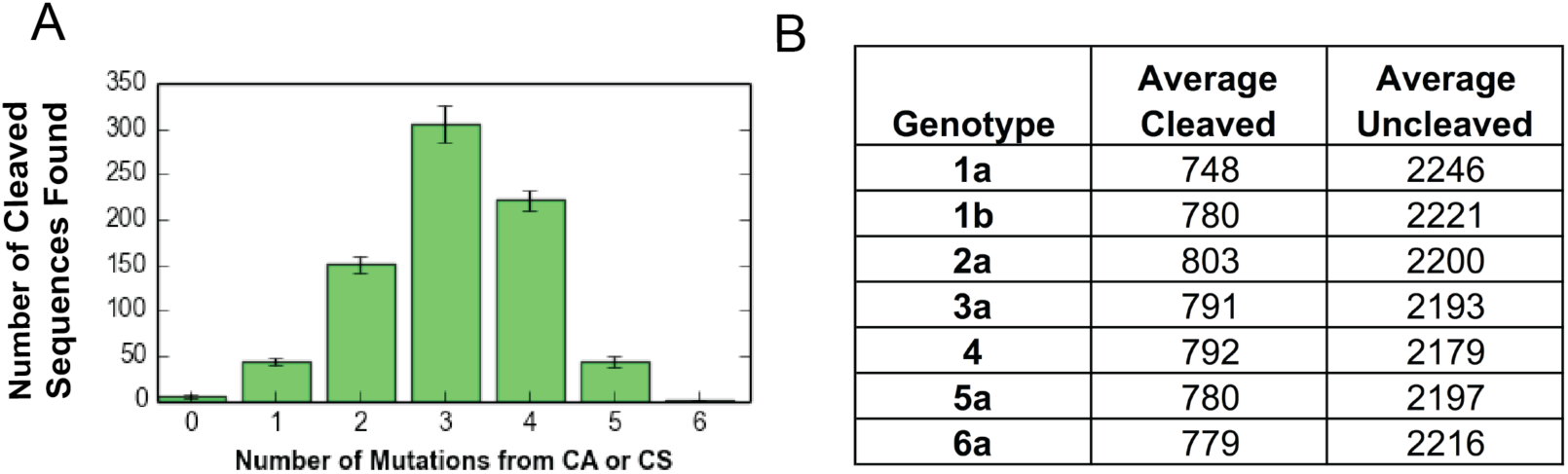
(A) Bar plot depicting the number of DNA mutations required to mutate from current protein sequence to ‘CS’ which is the scissile bond sequence for the HCV NS3/4A protease. (B) Table depicting the classification of all genotype derived 5-mers as classified by our SVM based predictor

### Specificity landscapes of Drug Resistant Protease variants

As the NS3/4A protease plays a key role in the viral assembly and maturation process, it is a target for therapeutics that aim at neutralizing viral activity. However, due to prevalence of quasispecies that are lurking at low levels in the population (Farci et al., 2000), several viral variants get exposed to the drug. Some of these develop resistance, and propagate to form Resistance Associated Variants (RAVs). To investigate how drug-resistant variants of the protease affect the mutational robustness, we explored the specificity landscape for three RAVs – A156T, D168A, R155K/A156T/D168A (Figure 7A). If the connectivity patterns of the sequences recognized by the RAVs are dramatically different (e.g., less clustered), it would indicate that their evolutionary fitness might be more limited under the heavy mutational load, as drifts on the substrate side would abolish the molecular interaction required for viral maturation. In this scenario, treatment with mutagens may be a desirable therapeutic strategy to induce error catastrophes. On the other hand, if similar mutationally robust connectivity is detected, the RAVs are likely to have a similar evolutionary potential as the wild type, and have an additional selective advantage in the population in the presence of the drug.

To obtain the landscapes of the protease variants (Supplementary Figure 5B-E), we generated the library using a PCR amplification based strategy; isolated functional variants using FACS, deep sequenced the isolated populations and validated mutants (Figure 7B, Supplementary Figure 2) identified from these populations using the yeast surface display assay. We find that the RAVs demonstrate a similar sequence profile to each other and to the wild type protease (Figure 7 C-F). Upon comparing the graphical properties of the specificity landscapes of the various protease variants, we observe that substrates that are experimentally detected in the cleaved pools of a greater number of protease variants are more reachable (higher pageranks) and more connected (higher degree) in each graph (Figure 7 G,H). As our goal was to compare gross features of the specificity landscapes for the wild type and variant proteases, we did not perform detailed structure-based calculations for RAVs. Nonetheless, these data indicate that more recognizable substrates appear to be more robust to changes in the protease, and indeed, mutational robustness is a key feature of this specificity landscape.

**Figure 7:**
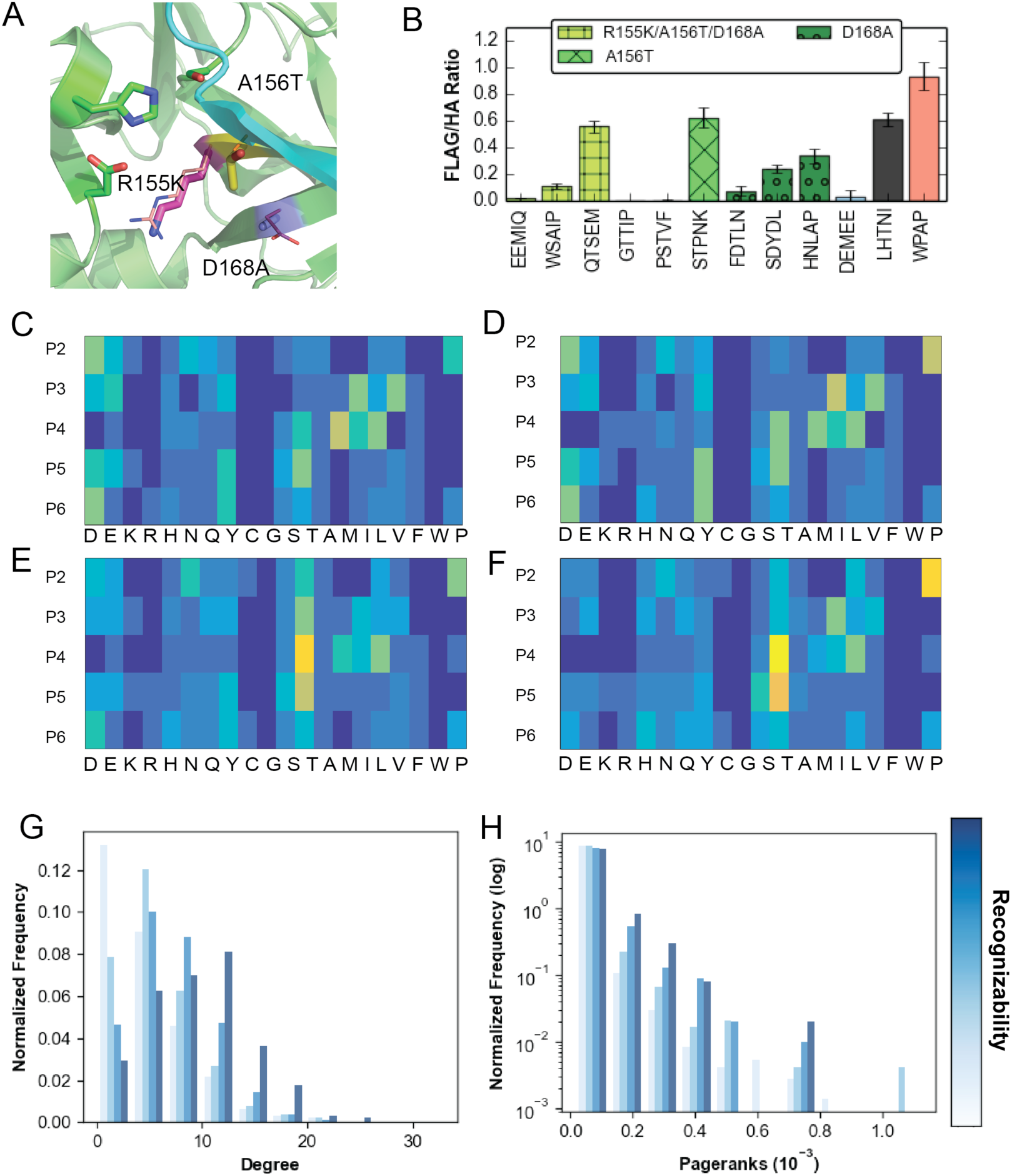
(A) Drug-resistant variant structures. Mutations are outlined in sticks and WT residues in lines. Active site residues are represented as green sticks. (B) Validation assay performed using yeast surface display for each of the mutants. (C-F) Mutant specificity logos for the triple mutant, D168A, A156T, and wild type showing that the mutants have very similar specificity profiles with slight variation as compared to the WT. (G-H) Substrate sequences that are recognized by a greater number of variants have higher degrees (G) and pageranks (H)

## Discussion

For RNA viruses, such as HCV, which have a high mutation rate, it has been hypothesized that viral evolution occurs *via* “survival of the flattest”: the most conserved viral form is not necessarily the most fit, but instead is the one most robust to mutation – thus mutational robustness may provide an evolutionary advantage (Lauring and Andino, 2010; Lauring et al., 2013; Wilke et al., 2001). Our data, based on combining, using a machine learning framework, information gleaned from library screening in yeast, deep sequencing, and structure-based modeling, provide atomic-resolution insight into how mutational robustness may be encoded in the molecular recognition landscapes involved in viral maturation, and indicate that cleavage specificity of the HCV NS3/4a protease is robust to patient-derived mutations in both the substrate regions as well as the protease. However, molecular interaction between the protease and substrate, which key for viral survival, is but one of the many evolutionary forces at play, especially in the “wild”(Boucher et al., 2016). Other factors such as the intrahost population size, stability and structure of the viral RNA genome, and interactions between the host and viral machineries and other environment dependent factors are also important to consider while considering evolutionary demands and trajectories.

We used a yeast surface display based assay that relies on the cleavage of the substrate region in the ER of yeast followed by cell sorting into gates and deep sequencing. We note that our assay is qualitative, and does not permit association the detected signal from deep sequencing to quantitative cleavability of substrates. Indeed, while we have validated that assignments to the three different pools is accurate with at least ~20 individual sequences, the identified cleaved and partially cleaved substrates may represent a wide range of catalytic efficiencies. A limitation of our technique is that it flattens this diversity into two pools. On the other hand, the assay construct with the protease and substrate on the same chain is a good representation of the situation in the virus, where the substrates of the protease are part of the same polyprotein (although both *cis* and *trans* cleavages occur) leading to high effective concentrations of substrates ([S] >> *K*_M_) *in vivo.* Under these saturating conditions in the virus and in our assay, we believe that selectivity and catalytic efficiency are both determined to a great extent by the goodness of fit of various substrates in the protease active site (i.e. by the *relative* binding between the different substrates). Similarly, our machine learning approach to combine experimental and computational data also is not without errors, showing a false-positive rate of ~5% on the experimental data. While we have validated several predictions on individual sequences (Figs. 1,3,7), it is possible that some individual sequences may be mispredicted. However, the overall trends regarding the connectivity patterns observed for the entire landscape should be robust to the misprediction noise. Further ongoing development of both computational and experimental methods we utilized is expected to help increase the accuracy of the approach.

HCV infects ~3% of the world population and the limited number of available viral genome sequences show low sequence heterogeneity in the protease and its substrate regions. Nevertheless, resistance mutations upon protease inhibitor drug treatment arise in a facile manner in the patient population, suggesting that genetic heterogeneity (quasispecies) indeed exists, possibly at levels too low for being captured in patient-derived sequencing. Spontaneous emergence of diverse HCV protease mutations was demonstrated recently by Liu and colleagues in continuous evolution studies of the protease (Dickinson et al., 2014), as well as by Sanjuan and colleagues in viral replicon assays coupled to ultradeep sequencing (Geller et al., 2016b). Our results show how genetic heterogeneity is entirely consistent with the properties of a key protease-peptide interaction in the virus, and therefore, provide a biophysical baseline for understanding evolvability of HCV, and evaluating inhibitor drug resistance risks. For example, our analysis suggests that viral evolution occurring at the substrate sites on the polyprotein could also contribute to drug resistance. Due to the flatness of the specificity landscape and high inter-connectedness of partially cleaved and fully cleaved clusters, novel sequences that are better substrates of drug-resistant variants may easily arise. Thus, considering both substrate and protease variation in evaluating and designing anti-viral therapies may be necessary. This mode of substrate coevolution-based drug resistance has been observed in HIV-1(Dam et al., 2009). At the same time, our analysis of the dominant HCV sequences obtained from patients suggests that the protease substrate interactions may also contribute to negative selection and help limit the acquisition of heterogeneity – the sequences of sites in the protease that are potentially cleavable upon acquisition of CA/CS at the P1-P1’ junction (Fig. 6) appear to be mutationally distant from doing so. Thus, the protease-substrate interaction landscape reveals that the balance between mutational robustness, negative selection and adaptive potential to environmental changes may be necessary to consider for understanding and therapeutic interventions.

In summary, our exploration of the specificity landscape uncovers novel specificities for the HCV NS3/4A protease and data provides a biophysical basis for the mutational robustness observed for a key interaction required in HCV propagation. Given the widespread prevalence of HCV, insights obtained here may help in better understanding, and tackling, the evolutionary trajectories of this ever-changing virus. The developed approach is general, and combining experimental deep sequencing and Rosetta-based structural modeling at a matching high throughput, followed by statistical machine learning, may be useful for elucidating a significantly larger space of sequence-function relationships for a variety of other systems.

## Author Contributions

SDK conceived the study. SDK, MAP, ABR designed experiments. MAP, ABR, DZ performed experiments. MAP, ABR, SDK analyzed the data. MAP, ABR, SDK wrote the paper.

## Acknowledgments

This work was supported by NSF grant MCB1330760 (to SDK) and NSF Graduate Research Fellowship Grant DGE-1433187 (to ABR). We thank L. Cuypers, H. Khiabanian. D. Kumar, T. Choi and S. Annavarappu for technical assistance, and T. Whitehead, A. Keating and D. Tawfik for helpful suggestions. We also thank Y. Li, B. Iverson, and G. Georgiou for sharing the LY104 plasmid used in the YESS assay experiments. This work used resources from the Rutgers Discovery Informatics Institute, which are supported by Rutgers and the State of New Jersey.

## Methods

### Two step screening approach to avoid stop codons

The LY104 vector was a gift from Y. Li, B. Iverson, and G. Georgiou (University of Texas at Austin). The library was constructed using a two-step screening approach to avoid enrichment of false positives. The first step was an expression screen, which was done by combining the library with a protease inactive vector (LY104 S139A knockout). The recombination was performed by homologous recombination technique in yeast EBY100 cells. We used a modified electroporation-based method as described by Kowalsky et al. The transformed library was allowed to grow for 48 hours at 30 C, up to an OD_600_ of 2.0. Dilutions of 1/10, 1/100 and 1/1000^th^ were plated from the initial culture to calculate the transformation efficiency and library size. The double positive cell population was isolated and enriched using a Fluorescence Assisted Cell Sorting technique. The expressible library was then recombined with a vector containing the active protease, using the aforementioned homologous recombination technique. This library of functional variants was allowed to grow up to 48 hours at 30 C and then sorted into three sequence pools – cleaved, partially cleaved and uncleaved. The gates for the FACS were defined using clonal substrates that displayed varying levels of cleavage activities. The three sequence pools were enriched via three rounds of successive selection (using FACS) and growth. The DNA from the three sequence pools was extracted using the Omega E.Z.N.A yeast plasmid kit. Biological duplicates were sequenced to get an estimate of error correction necessary for post processing this data.

### Library Generation methodology

The library was constructed using a PCR amplification based technique using NNK mixed base oligonucleotides (Integrated DNA Technologies). The LY104 vector was linearized using DNA oligonucleotides (IDT). The NNK library insert (~576 bp) and linearized vector (~6000 bp) were combined using Homologous Recombination using electro-competent EBY100 yeast cells. The transformed EBY100 cells were rescued using a YPD medium and allowed to grow in a 250 μL Selective Complete Growth Medium (-UW). The media was supplemented with 250 μL of Ampicillin and Kanamycin to avoid bacterial contamination.

### Library Testing and Enrichment

The transformed library was allowed to grow for ~48 hrs (upto OD_600_ 2.0) and then induced and tested using Flow cytometry. 1.5 x 10^7^cells(OD_600_ ~0.5) were pelleted and resuspended in 2 mL induction media (20g/L galactose, 2 g/L glucose) supplemented with 2 μL each of 1000x antibiotics (carbenicillin, kanamycin). The induction cultures were grown overnight at 30 C (225 rpm) to an OD_600_ of 1-1.5. All spins in the protocol were done at 3000 r.c.f for 5 min. The induced cultures were pelleted and washed with 500 μL PBS followed by 500 μL PBS+ 0.5% BSA. 1 μL of each antibody stain (anti-FLAG PE from Prozyme, PJ315 and anti-HA FITC from Genscript, A01621) was incubated with 10^7^ cells for 30 min at 4 C. The samples were resuspended by vortexing and incubated at RT for an additional 30 min. The cells were washed with 100μL PBS with 0.5% BSA, pelleted and then resuspended in 500 μL PBS. Samples were diluted to achieve a final concentration of 10^6^ cells/mL and then FITC (anti-HA) and PE (anti-FLAG) intensities were detected using a Flow Cytometer (Beckman Coulter Gallios).

The tested cells were then enriched using a MoFlo XDP Cell Sorter (final cell density 10^7^). Up to 10^6^ cells were collected and grown in the Selective Complete Growth Media for 48 Hours. Two rounds of sorting and enrichment were carried out to select for clones that were expressed. The selected cells were grown for 48 hours. The DNA from the selected cell population was extracted using E.Z.N.A Zymoprep Kit (Omega).

### Cell Sorting into Cleaved, Uncleaved, Partially Cleaved Populations

The expressible fraction of the library was combined with the active LY104 vector using a second round of Homologous recombination following the same protocol as mentioned above. Using the MoFlo XDP Cell Sorter we defined Cleaved, Uncleaved and Partially cleaved gates for further selection of this population. These gates were defined based on previously experimentally tested sequences.

These cells from the selected population were put through three rounds of enrichment and sorting. In the first round of sorting, cells were collected into two gates – Cleaved and Uncleaved. The Uncleaved sample was further enriched in the second sorting round whereas the Cleaved population was separated into Cleaved and Partially cleaved gates. Cells were collected for each sorting round until a cell count of 10^6^ was reached. At the culmination of each sorting round, DNA was collected from each population by using a Zymoprep Kit (Omega).

### Preparation for Illumina Sequencing Run

The DNA samples collected from each of the populations were prepared by 25 cycle amplification (Kowalsky et al. 2015) with inner primers (Supplementary Table 3). The samples were then run on a 1% Agarose gel to confirm the amplification of a single species. These were further amplified using 8 PCR cycles to include the DNA barcode used in the deep sequencing protocol and checked for quality using a Bioanalyzer 2100. The Deep sequencing was performed on a NextSeq 500 (Illumina) giving a 75 bp paired end read.

### Expression Protocols

#### Protease expression

Expression and purification protocol was a modification of previously published protocols (Wittekind et al. 2001; Gallinari et al. 1998; Romano et al. 2012). Transformed BL21 (DE3) *E. coli* cells were grown at 37°C and induced at an optical density of 0.6 by adding 1 mM IPTG. Cells were harvested after 5 hours of expression, pelleted, and frozen at –80°C for storage. Cell pellets were thawed, resuspended in 5 mL/g of resuspension buffer (50 mM phosphate buffer, 500 mM NaCl, 10% glycerol, 30 mM imidazole, 2 mM β-ME, pH 7.5) and lysed with a sonicator. The soluble fraction was retained, applied to a nickel column (Qiagen), washed with resuspension buffer, and eluted with resuspension buffer supplemented with 200 mM imidazole. The eluent was dialyzed overnight (MWCO 10 kD) into a protease storage buffer (20mM Tris.HCl,pH 8.0, Glycerol 20%, 100 mM KCl, 1mM DTT, 0.2 mM EDTA) to remove the imidazole. The purified protein was then flash frozen and stored at -80 C.

#### Substrate (MBP-GST construct) expression

The transformed BL21(DE3) cells were grown at 37 C to an optical density of 0.6 and induced using 0.2 mM IPTG. Upon induction the cells were grown overnight at 18 C. the cells were harvested and the cell pellet was resuspended in a resuspension buffer (50 mM Tris.HCl, pH8.0, 500 mM NaCl, 30 mM immidazole). The resuspended cells were lysed via sonication and the soluble fraction was applied to a Nickel column (Qiagen). The column was washed using the resuspension buffer and then the protein eluted using an Elution buffer-50 mM Tris.HCl, pH8.0, 150 mM NaCl, 300 mM imidazole. The protein was dialyzed overnight to remove the imidazole and frozen until use.

### Gel based validation assay

The frozen aliquots of substrate solutions were thawed and dialyzed overnight into the reaction buffer (50mM HEPES (pH 7.5), 150 mM NaCl, 0.1% Triton X-100, 15% Glyecerol, 10mM DTT). 28.5 nM substrate was incubated overnight with 500nM, 700nM, 1uM, 2uM, 3uM and 4 uM protease. The resultant reactions were run on a SDS PAGE gel to check for cleavage activity.

## Sequence Processing

### Sequence Alignment and Trimming

Data was received oriented in the correct orientation and filtered for quality of 20. Each sequence was searched for the presence and location of “TCTTTATAA”, a unique string within the WT sequence, to align the sequences. If the index of “TCTTTATAA” in sequence *a* is less than the index of “TCTTTATAA” in the WT sequence, the beginning of sequence *a* is padded to match the beginning of the WT sequence. If the index of “TCTTTATAA” in sequence *a* is greater than the index of “TCTTTATAA” in the WT sequence, the beginning of sequence *a* is truncated to match the beginning of the WT sequence. If “TCTTTATAA” is not found in sequence *a*, it is discarded. If the padded or truncated sequence *a* is shorter than the index of the library region in the WT sequence, sequence *a* is discarded. If sequence *a* is longer than the index of the library region but shorter than the WT sequence, the end of sequence *a* is padded to match the WT sequence. Finally, we check that the padded or truncated sequence *a* matches the WT sequence entirely except for the library region. If it does not match the WT sequence, we discard sequence *a.*

### Threshold Determination

After aligning and trimming sequences, we calculate a normalized count of each sequence so that the sum of the normalized counts in each population is equal. This is achieved by multiplying each sequence count in population *a* by a normalization factor that is equal to the number of sequences in the largest library divided by the number of sequences in library *a*. Then, to determine the minimum frequency of each sequence in the population above which we are confident of the validity of its representation in the library, we used several methods:

#### 1) Overlap between cleaved and uncleaved sequences

We expect little overlap between the populations of cleaved and uncleaved sequences. However, at low counts, there is some overlap between the two populations. For each threshold, we calculated the number of sequences that overlapped between cleaved and uncleaved sequences, and normalized by the count of unique translated cleaved DNA sequences at that threshold. We determined the amount of overlap as a percentage of the initial overlap between the populations at a threshold of 1, and then found the threshold that gave >= 10% of the initial overlap (see Supplementary Figure 1). We repeated this analysis for all four variant populations. The threshold was less than or slightly greater than 11 for all variants.

#### 2) Duplicate population error

We sampled biological duplicates for the third round of enrichment for cleaved, uncleaved and partially cleaved sequence pools. As a post - processing step in the pipeline, we introduced duplicate population error correction, by plotting the difference in counts for common sequences of the biological duplicate samples and plotting against the counts in the first sample.

#### 3) SVM Convergence

In order to select for the threshold that gave us the most distinct populations, we generated cleaved and uncleaved sequence sets for thresholds 5,10,11,12,13,14, 15, 16, 25, 50, 75 and 100. Using an SVM based technique described previously (JMB) we calculated the auROC for all cleaved and uncleaved sequence populations for the listed thresholds. This enabled us to identify which threshold increases the distinction between the two populations.

**We decided upon a frequency threshold of 11 as one that satisfies all categories of threshold determination.**

### Enrich Software

We used a modified version of the Enrich software (Fowler et al. 2010) to find an enrichment ratio (ER) for each sequence. We only included sequences that had a normalized count (as defined above) of greater than or equal to eleven for both the unselected and selected populations. The enrichment ratio of sequence *v* in population *X* is defined using Equation 1. *F*_*v*,*X*_ is the frequency of sequence *v* in population *X*.

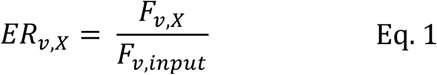

### Population Categorization

Sequences were sorted into one of three pools (cleaved, uncleaved and middle), based on the following criteria. Sequences that had a positive ER for more than one pool were discarded. Sequences that had a positive ER for either or both replicates for one pool only were assigned to that pool. Negative ERs were ignored.

We also sorted a second set with more stringent criteria, which was then used for training the SVM. For this set, if a sequence was found in more than one pool (even if it had a negative ER in the second pool), it was discarded. Additionally, only sequences with a positive ER > 2.0 were considered.

## Computational

### Graph Generation

Graph generation was done using Gephi 0.9.1 (Bastian et al. 2009). Nodes were assigned a fitness of 2.0 for cleaved nodes, 1.5 for middle nodes, and 1.0 for uncleaved nodes. Edge directionality was determined by distance from DEMEE, the starting sequence for library generation; in the case of edge *a* connecting nodes *b* and *c*, the node with a smaller hamming distance from DEMEE was chosen as the source for edge *a*. Edge weight was defined as the ratio of the starting sequence fitness to the ending sequence fitness. The graph layout was run in two steps, starting with a Fruchterman-Reingold layout to separate the nodes and then ending with the ForceAtlas2 layout to generate a force-directed graph. All statistics were run with Gephi default settings.

### Random Graph

The edges in the wild-type HCV graph were randomized using the following process. The source of each edge was kept and a population (cleaved, middle, or uncleaved) was randomly chosen for the target of the edge. The target of the edge was then randomly chosen from among that population.

### SVM Sequence Features

We used an encoding scheme that included twenty binary features per amino acid residue, where one of those features was a one and the rest were zeroes. The placement of the one was dependent on the identity of the amino acid. With five amino acid residues per sequence, this resulted in 100 total sequence features.

### Mutual Information

Correlation between residues at different positions was calculated using a mutual-information based metric (Equation 2), with modifications based on Buslje et al. (Equation 3) (Buslje et al. 2009) and Gouveia-Oliveira and Pedersen (Equation 4) (Gouveia-Oliveira & Pedersen 2007). We begin with MI between amino acid *a* at position *i* and amino acid *b* at position *j* defined as:

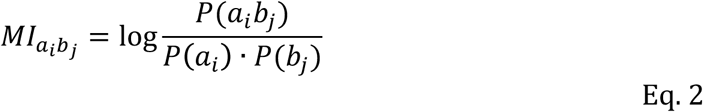

*P*(*a_i_*) and *P*(*a_i_b_j_*) are defined with a pseudocount to correct for MSAs with low counts.

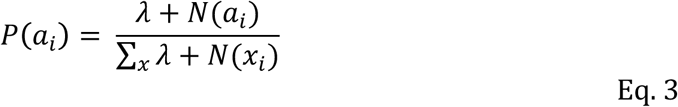

*N*(*a_i_*) is the count of amino acid *a* appearing at position *i*. *λ* is equal to the length of sequences in the MSA divided by 20 for single-amino acid counts (*N*(*a_i_*)) and 400 for double-amino acid counts *N*(*a_i_b_j_*). We also modified MI to include row-column weighting:

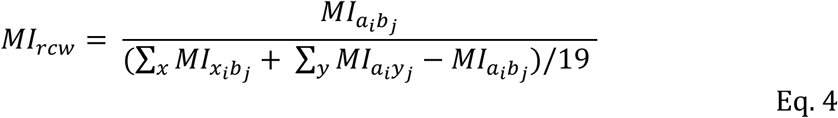

#### Obtaining viral genomes from patient populations

The list of complete viral polyprotein genomes was accessed and downloaded from NCBI. These genomes were checked to ensure that the sequence covered all NS3 substrate regions. We translated the DNA sequence that we downloaded from NCBI into a protein sequence and compared the five substrate regions “DLEWTST”, “DEMEECASHL”, “EDVVCCSM”, ECTTPCSGS” and “ALVTPCASH” to discover the diversity found in the substrate region for the patient genomes.

The dataset of aligned genomes utilized in Cuypers et al. was used for dN/dS measurements and for the mapping of predicted cleaved and uncleaved sequences within the genome (Cuypers et al. 2015).

## Supplementary Figures

**Supplementary Figure 1:**
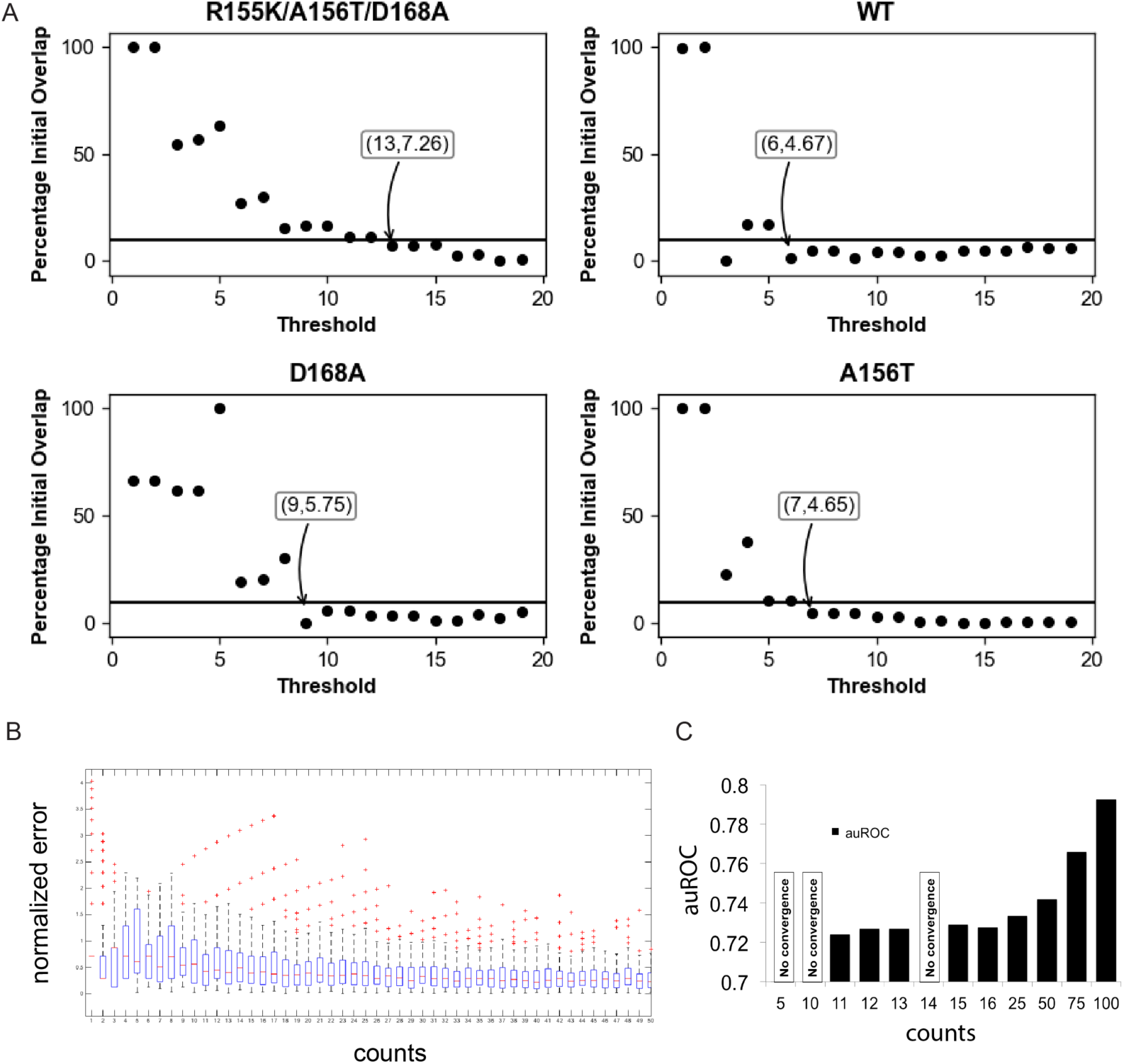
Threshold Determination. (A) Threshold vs. percentage of initial overlap between cleaved and uncleaved sequences for all 5 variants. The final threshold beyond which all other thresholds have a percentage overlap that is <= 10% is marked with an arrow. (B) Duplicate population analysis. Normalized error is calculated for biological duplicates of cleaved samples, by the formula: |(counts_S2-counts_S)| / counts_S2, where sample S and S2 are biological duplicates. (C) The Area Under the Curve for the ROC plot, when the SVM is used to classify cleaved versus uncleaved sequence pools at various count thresholds.

**Supplementary Figure 2:**
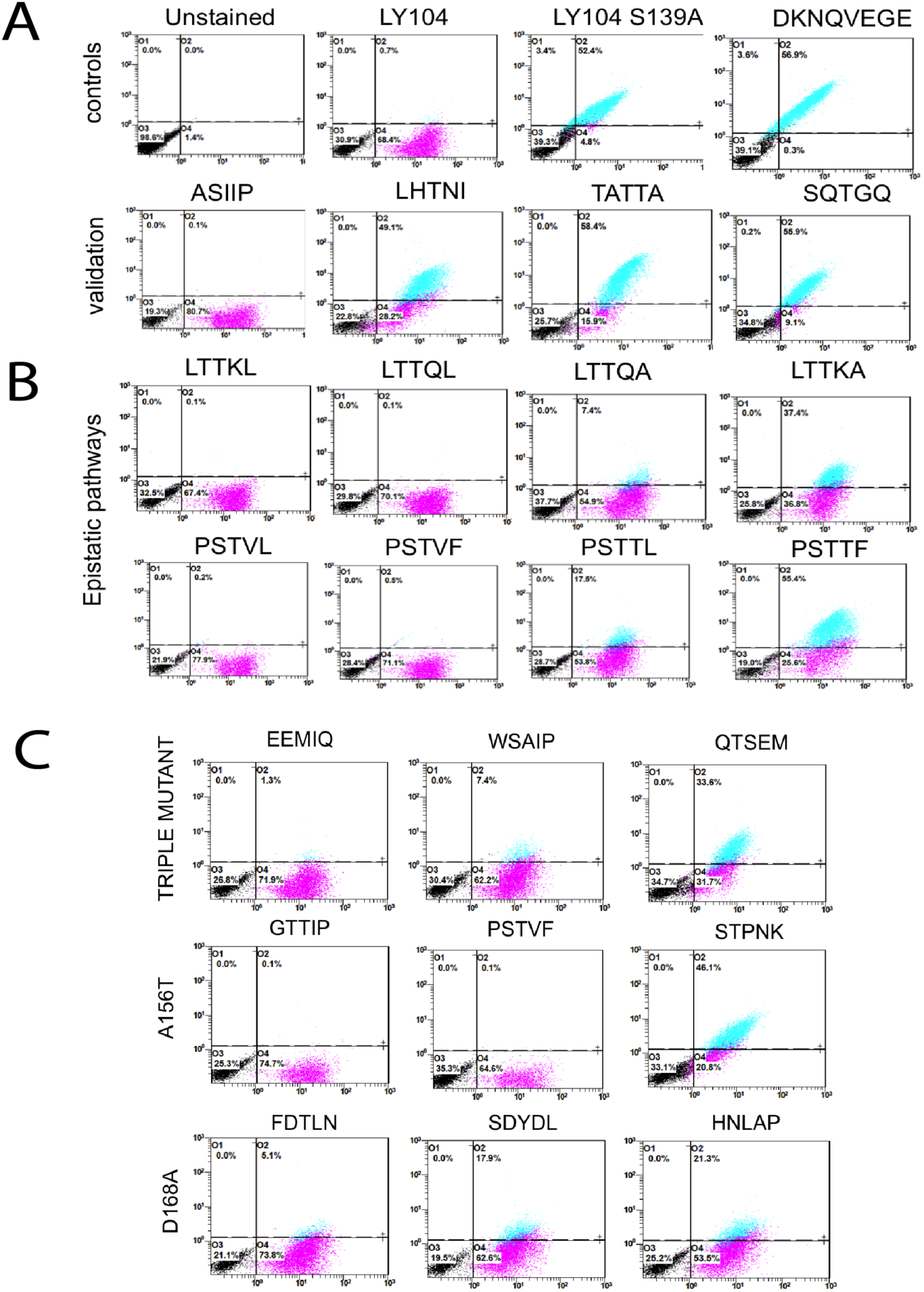
The figure displays 2D plots of anti HA and anti FLAG antibody signals seen in the Flow cytometry assay. (A) displays controls (B) Epistatic pathway validation (C) Drug resistant mutant validation plots

**Supplementary Figure 3:**
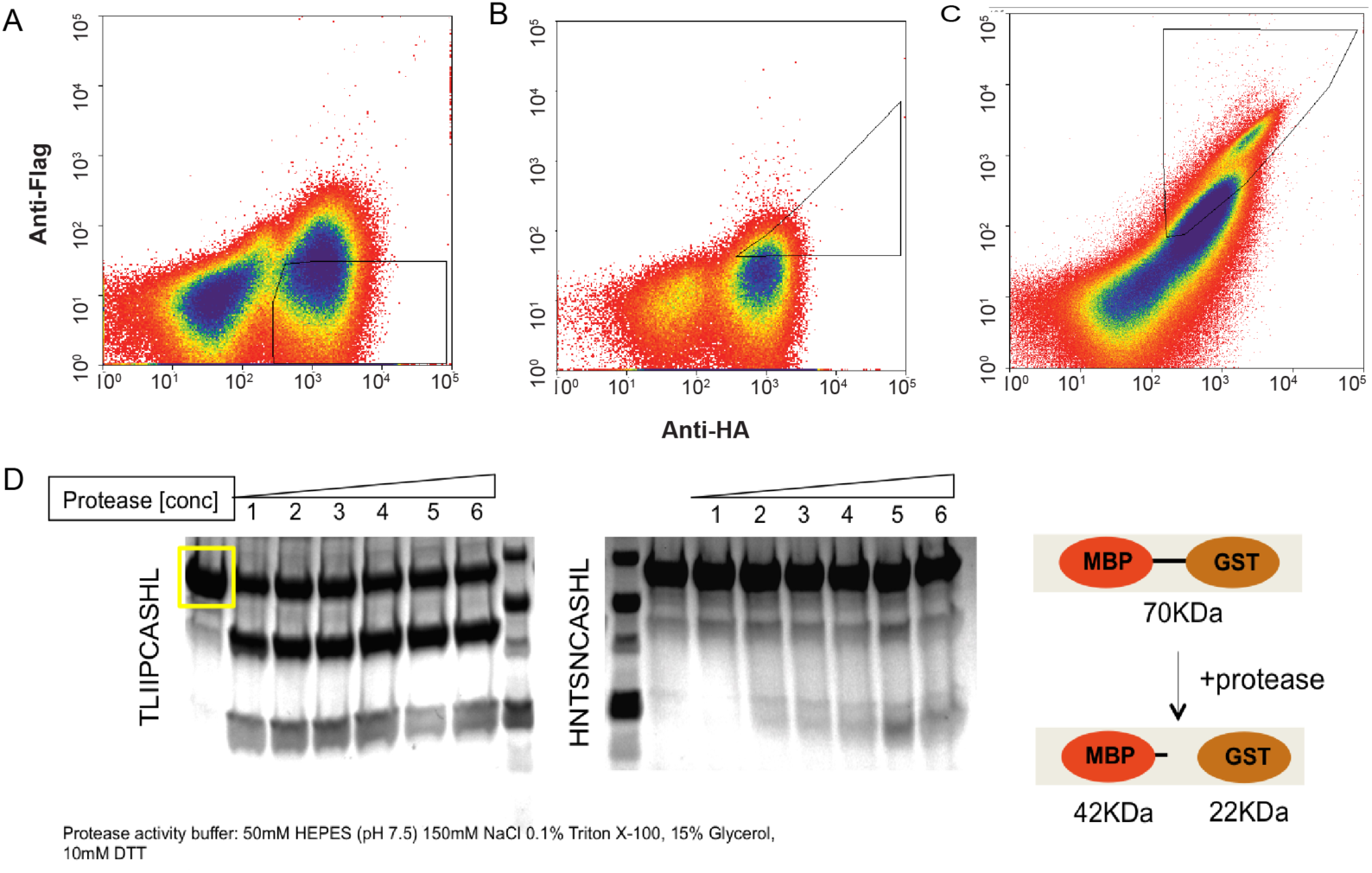
Flow cytometry 2D plots showing anti-HA and anti -FLAG stains for cell populations collected after enrichment round three. (A) Plot showing gate and cell population for cleaved (B) partially cleaved and (C) uncleaved populations. (D) In vitro gel based assay using an MBP -GST fusion protein (70KDa). Upon overnight incubation with increasing concentrations of the protease-500nM, 700 nM, 1uM, 2uM, 3uM, 4uM (well 1 through 6) results in cleavage for substrate TLIIPCASHL whereas HNTSNCASHL displays no cleavage.

**Supplementary Figure 4:**
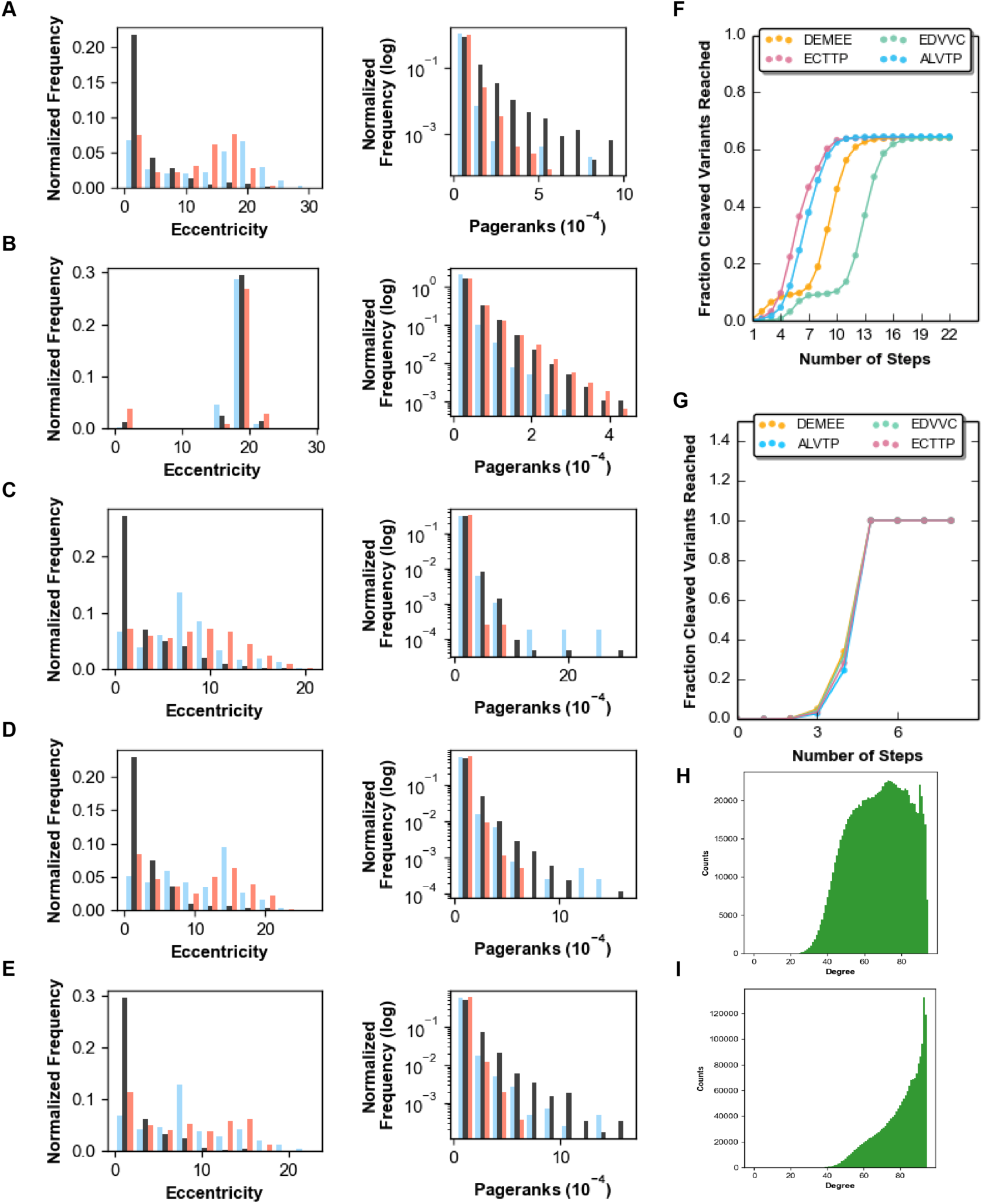
Cleaved (blue), uncleaved (red), and partially cleaved (black) graph metrics for (A) wildtype HCV (B) randomly generated graph (C) R155K/A156T/D168A triple mutant (D)A156T (E) D168A. Partially cleaved sequences generally have higher pageranks and lower eccentricity. Number of mutations vs. fraction cleaved variants reached for (F) experimental and (G) SVM-generated graphs. (H) degree distribution for cleaved sequences in SVM derived graph (I) degree distribution for uncleaved seqs

**Supplementary Figure 5:**
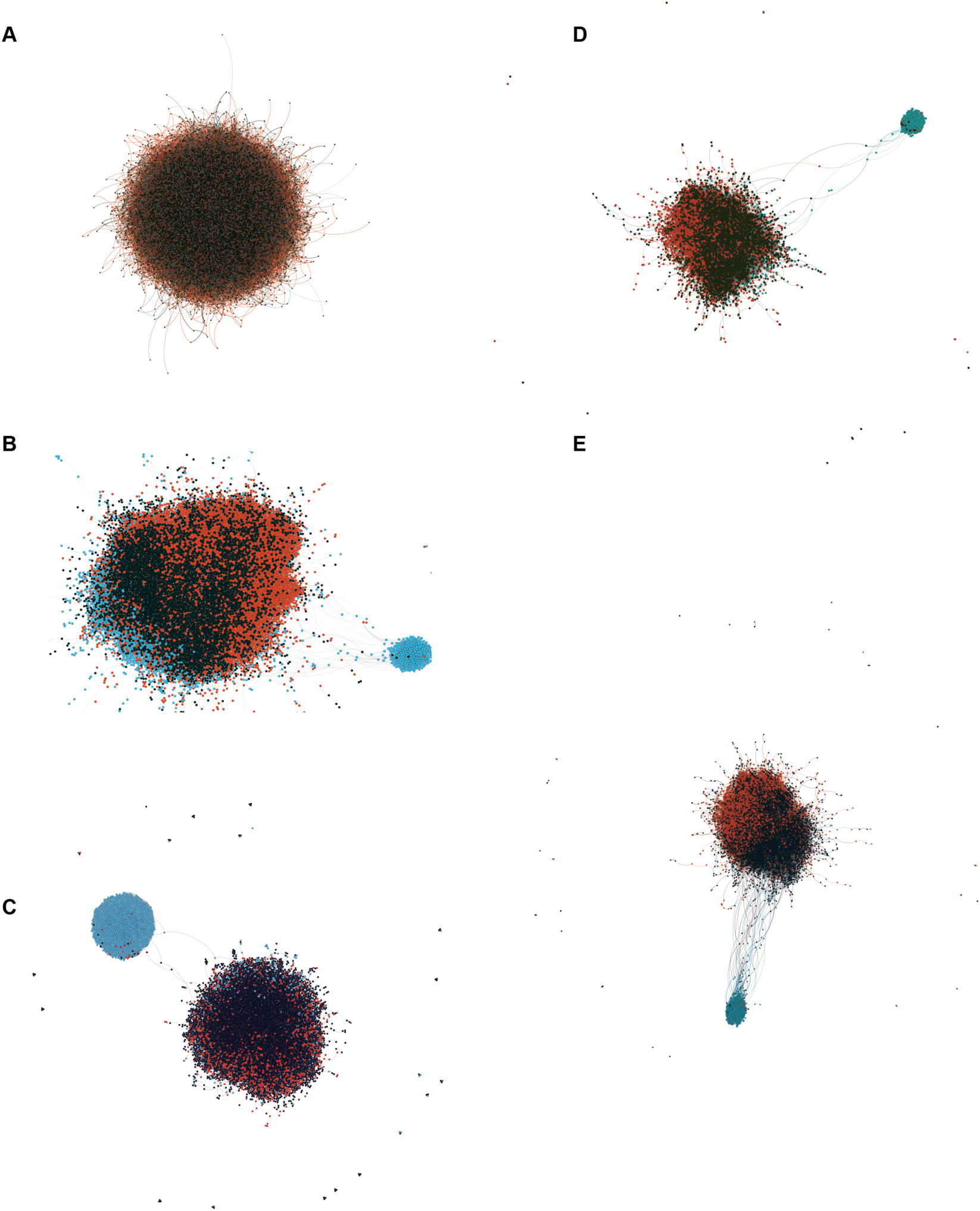
Force-directed graphs for (A) randomly generated graph (B) wild-type HCV protease (C) R155K/A156T/D168A triple mutant (D) D168A variant (E) A156T mutant

**Supplementary Figure 6:**
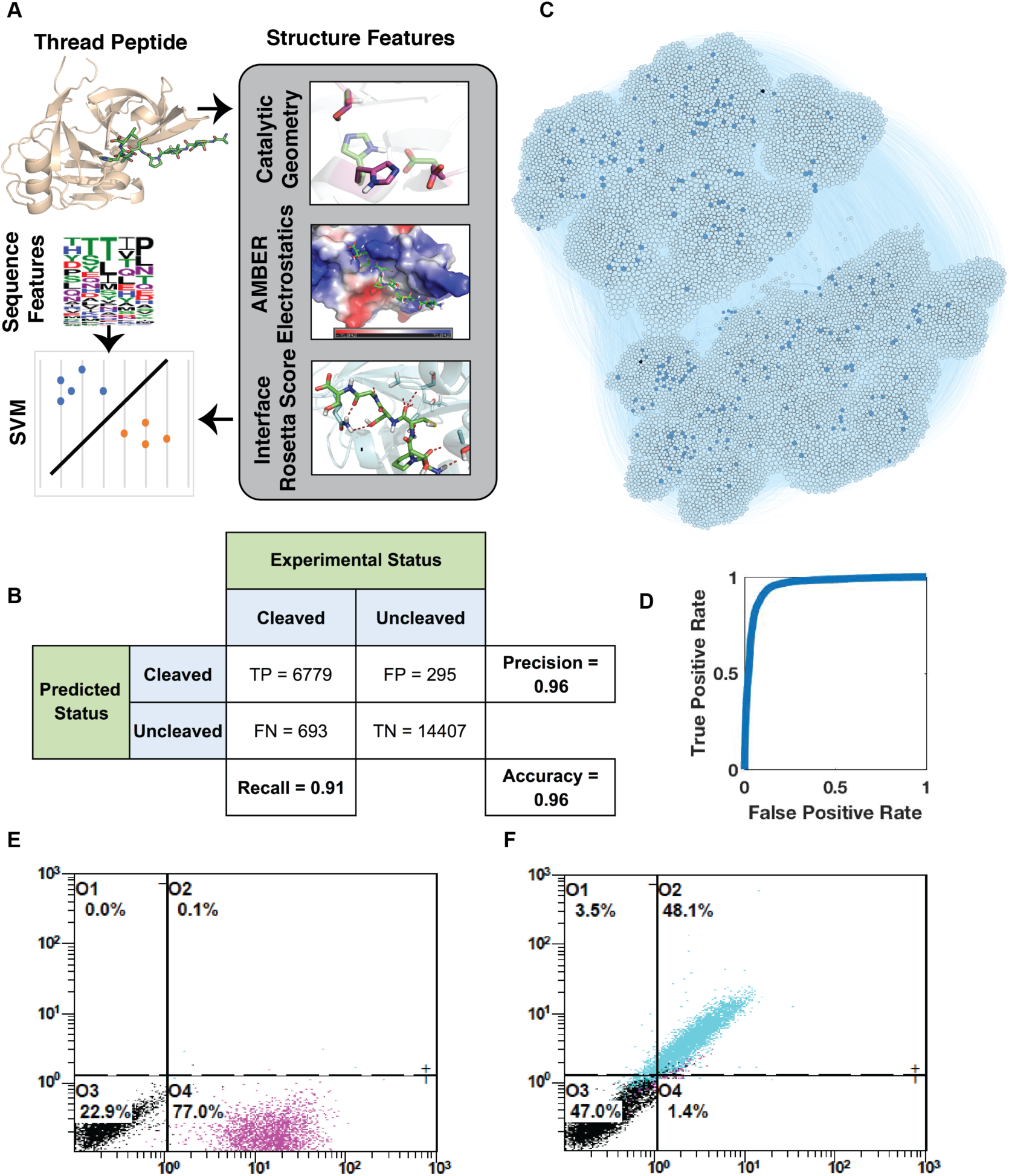
SVM predictions. (A) Schematic workflow for SVM generation. (B) Sub-graph of SVM predicted cleaved sequences with a distance > 2 from the hyperplane. Experimental cleaved sequences are dark blue and experimental partially cleaved sequences are black. (C) Contingency table for SVM prediction. (D) ROC plot of cross-validation on training set for SVM. (E) Flow cytometry plot for ECTIP (SVM-predicted cleaved). (F) Flow cytometry plot for RPGPG (SVM-predicted uncleaved).

**Supplementary Figure 7:**
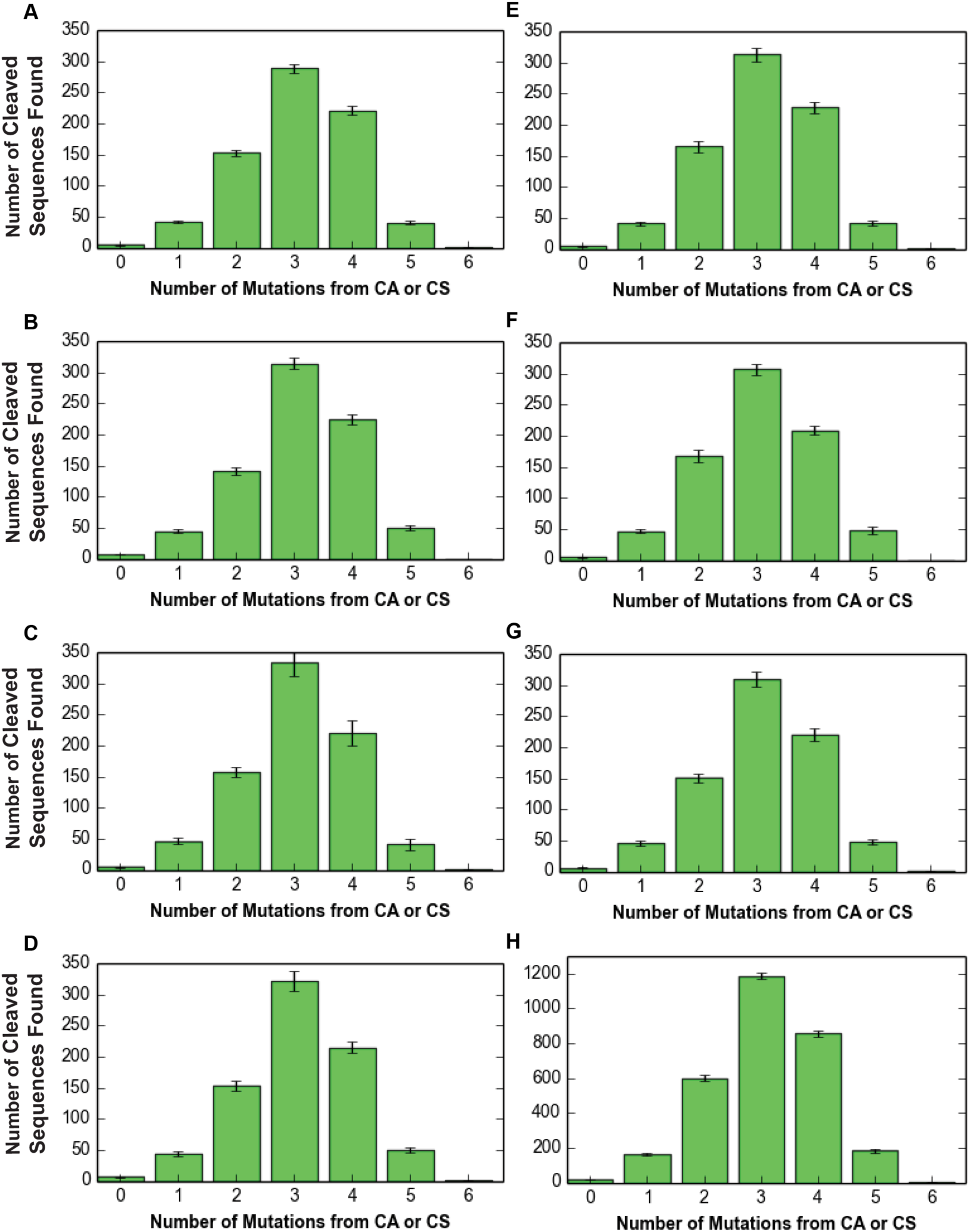
Bar plot depicting the number of DNA mutations required to mutate from current protein sequence to ‘CS’ which is the scissile bond sequence for the HCV NS3/4A protease for (A) strain 1a, (B) strain 1b, (C) strain 2, (D) strain 3, (E) strain 4, (F) strain 5, (G) strain 6, (H) control. Control is the distance from CA/CS for all 2-mers in all genotypes.

## Supplementary Tables

**1.**
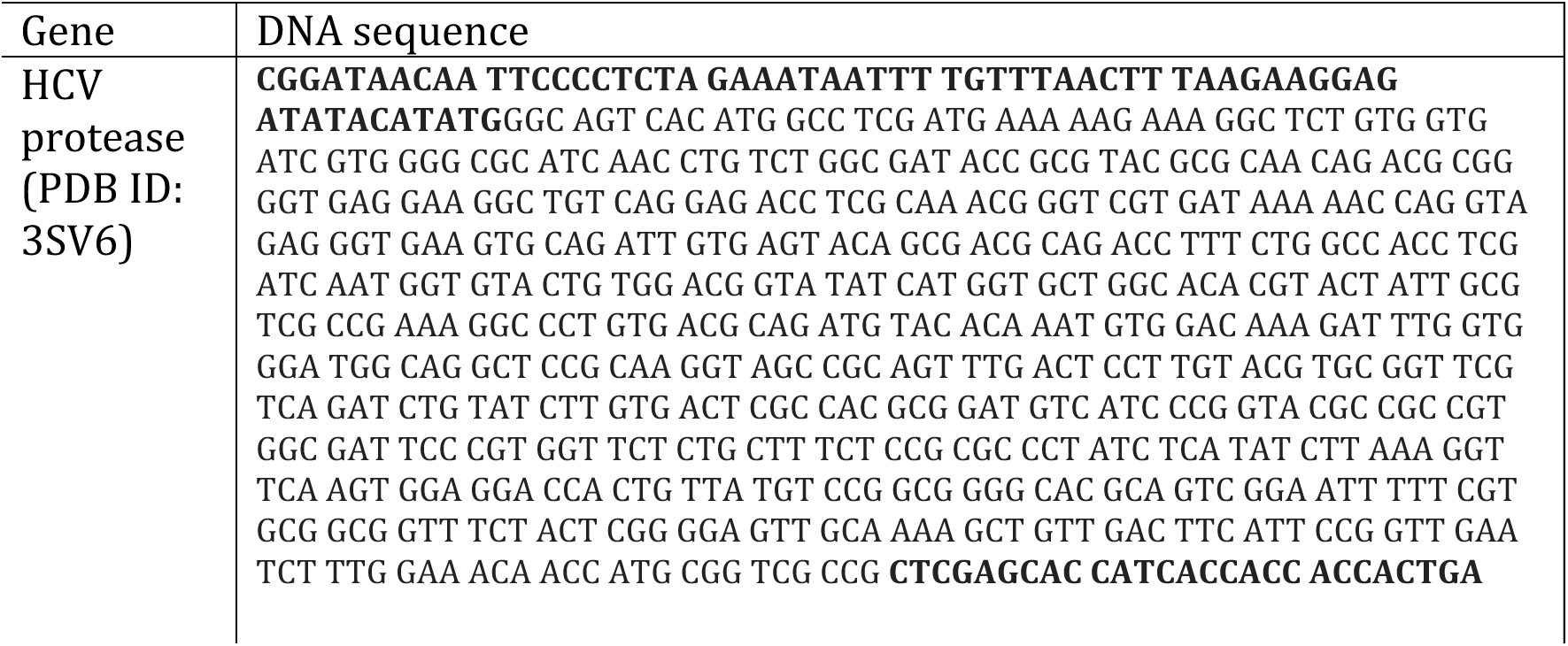
Genes.

**2.** Cell sorting statistics.

2. Cell sorting statistics:
|  | Functional pool | Sort Round | Cell # |
| --- | --- | --- | --- |
| 1 | CLEAVED<br>UNCLEAVED | 1 | 420 K<br>109 K |
|  | CLEAVED<br>MIDDLE<br>UNCLEAVED | 2 | 1.05M<br>105K<br>775K + 295K |
|  | CLEAVED<br>MIDDLE<br>UNCLEAVED | 3 | 1.55M<br>89K<br>675K |
| 2 | CLEAVED<br>UNCLEAVED | 1 | 1 M<br>205 K |
|  | CLEAVED<br>MIDDLE<br>UNCLEAVED | 2 | 1.15M<br>300K<br>1.05 M |
|  | CLEAVED<br>MIDDLE<br>UNCLEAVED | 3 | 2M<br>262K<br>707K |
| 9 | CLEAVED<br>UNCLEAVED | 1 | 812K + 2.65 M<br>359K |
|  | CLEAVED<br>MIDDLE<br>UNCLEAVED | 2 | 1.4 M<br>94 K<br>1.02 M |
|  | CLEAVED<br>MIDDLE | 3 | 1.77 M<br>324 K |
|  | UNCLEAVED |  | 1.5 M |
| 10 | CLEAVED<br>UNCLEAVED | 1 | 2.7 M<br>646 K |
|  | CLEAVED<br>MIDDLE<br>UNCLEAVED | 2 | 1.04M<br>183K<br>1.06 M |
|  | CLEAVED<br>MIDDLE<br>UNCLEAVED | 3 | 1.59M<br>1.16M<br>1.5M |

**3.**
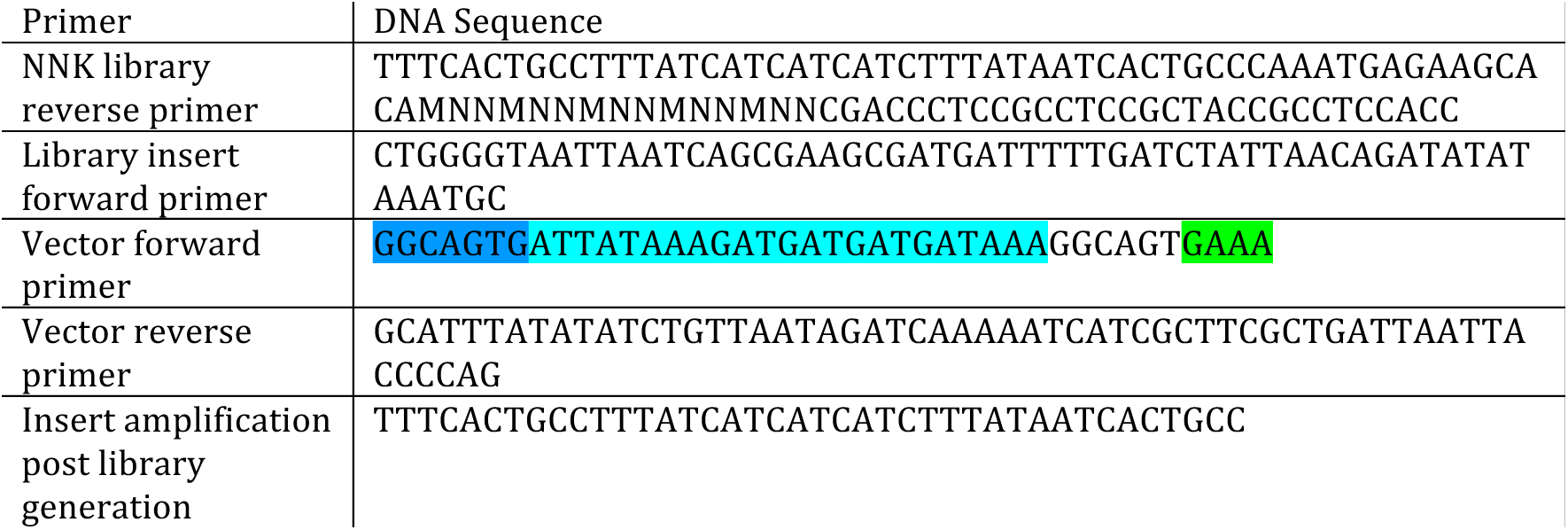
List of oligos for next - sequencing library generation.

**4.**
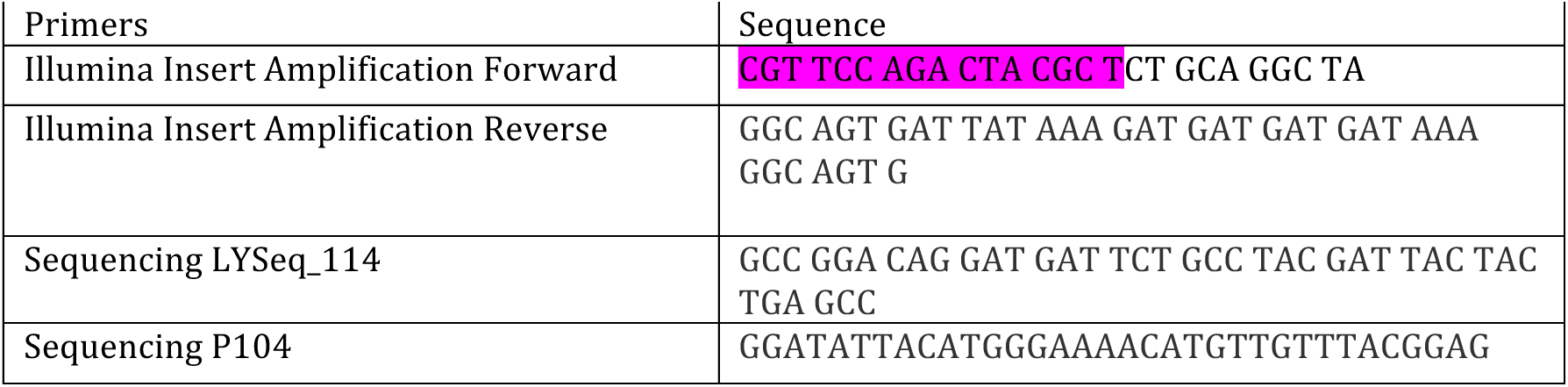
List of oligos for Illumina sample prep and sequencing.

**5.** Deep sequencing processing statistics.

5. Deep sequencing processing statistics
| Variant | Population | Initial | Post-thresholding | Post-categorization |
| --- | --- | --- | --- | --- |

|  |  | Unique Counts | Unique Ratios | Unique Counts | Unique Ratios | Unique Sequences |
| --- | --- | --- | --- | --- | --- | --- |
| WT | Background | 379361 |  | 74575 |  |  |
|  | Cleaved-Rep1 | 216254 | 84773 | 30328 | 23550 | 7472 |
|  | Cleaved-Rep2 | 260764 | 95730 | 29238 | 23692 |  |
|  | Partial-Rep1 | 219369 | 89830 | 32355 | 22690 | 8737 |
|  | Partial-Rep2 | 354253 | 123573 | 28631 | 21986 |  |
|  | Uncleaved-Rep1 | 587740 | 183536 | 39235 | 32298 | 14702 |
|  | Uncleaved-Rep2 | 473115 | 160980 | 39115 | 32053 |  |
| R155K/<br>A156T/<br>D168A | Background | 339049 |  | 64406 |  |  |
|  | Cleaved | 139722 | 50895 | 16374 | 10948 | 3135 |
|  | Partial | 270663 | 108408 | 40602 | 29624 | 11562 |
|  | Uncleaved | 209209 | 75869 | 23432 | 10425 | 3703 |
| A156T | Background | 367896 |  | 68199 |  |  |
|  | Cleaved | 140479 | 52199 | 18718 | 9911 | 3644 |
|  | Partial | 251274 | 95066 | 26348 | 17151 | 8461 |
|  | Uncleaved | 277994 | 109684 | 29935 | 17594 | 9564 |
| D168A | Background | 314942 |  | 65787 |  |  |
|  | Cleaved | 197578 | 65957 | 19018 | 10348 | 4350 |
|  | Partial | 336654 | 108567 | 30535 | 16929 | 5780 |
|  | Uncleaved | 286784 | 96578 | 26993 | 15155 | 7514 |

**6.**
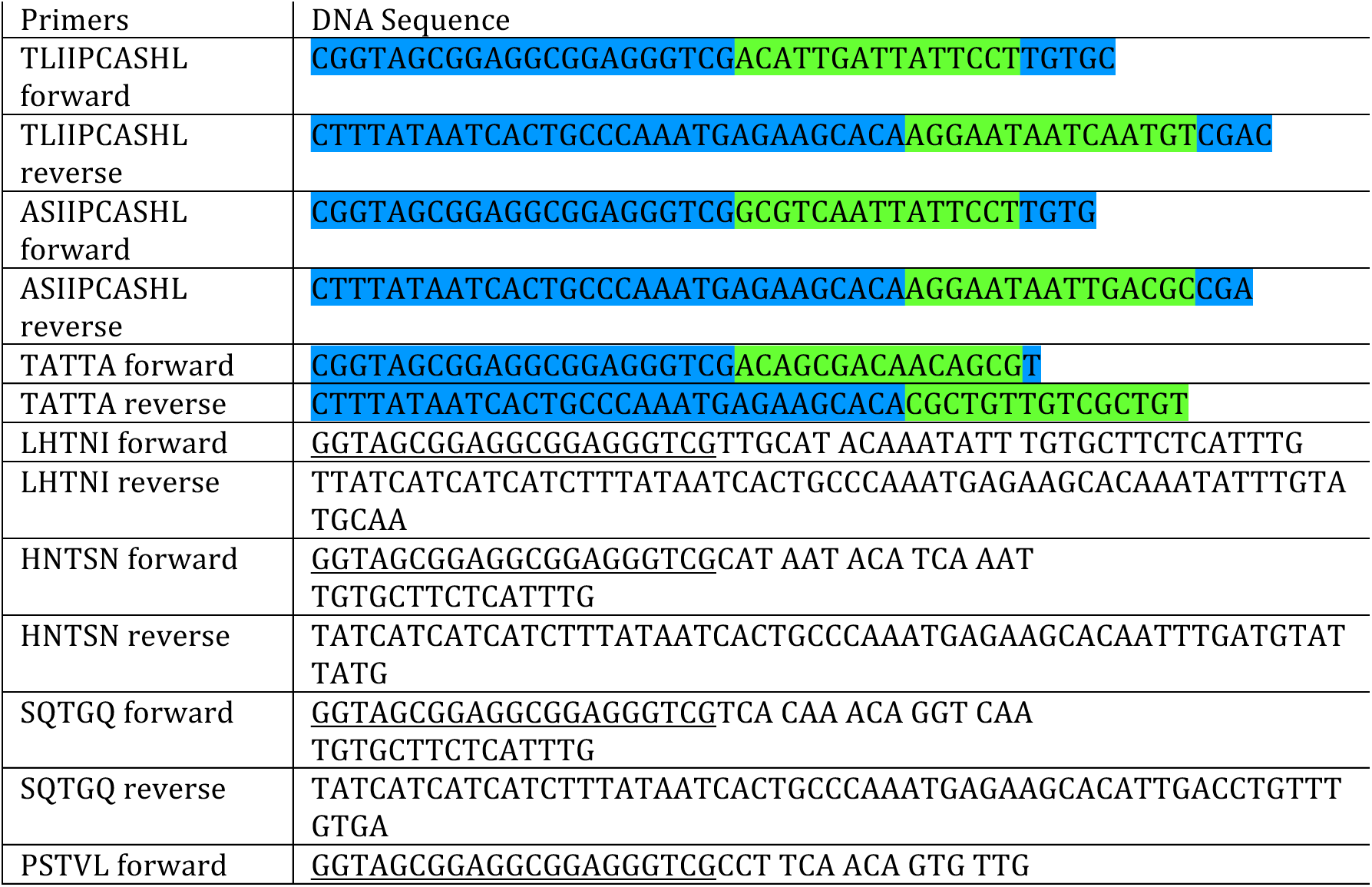
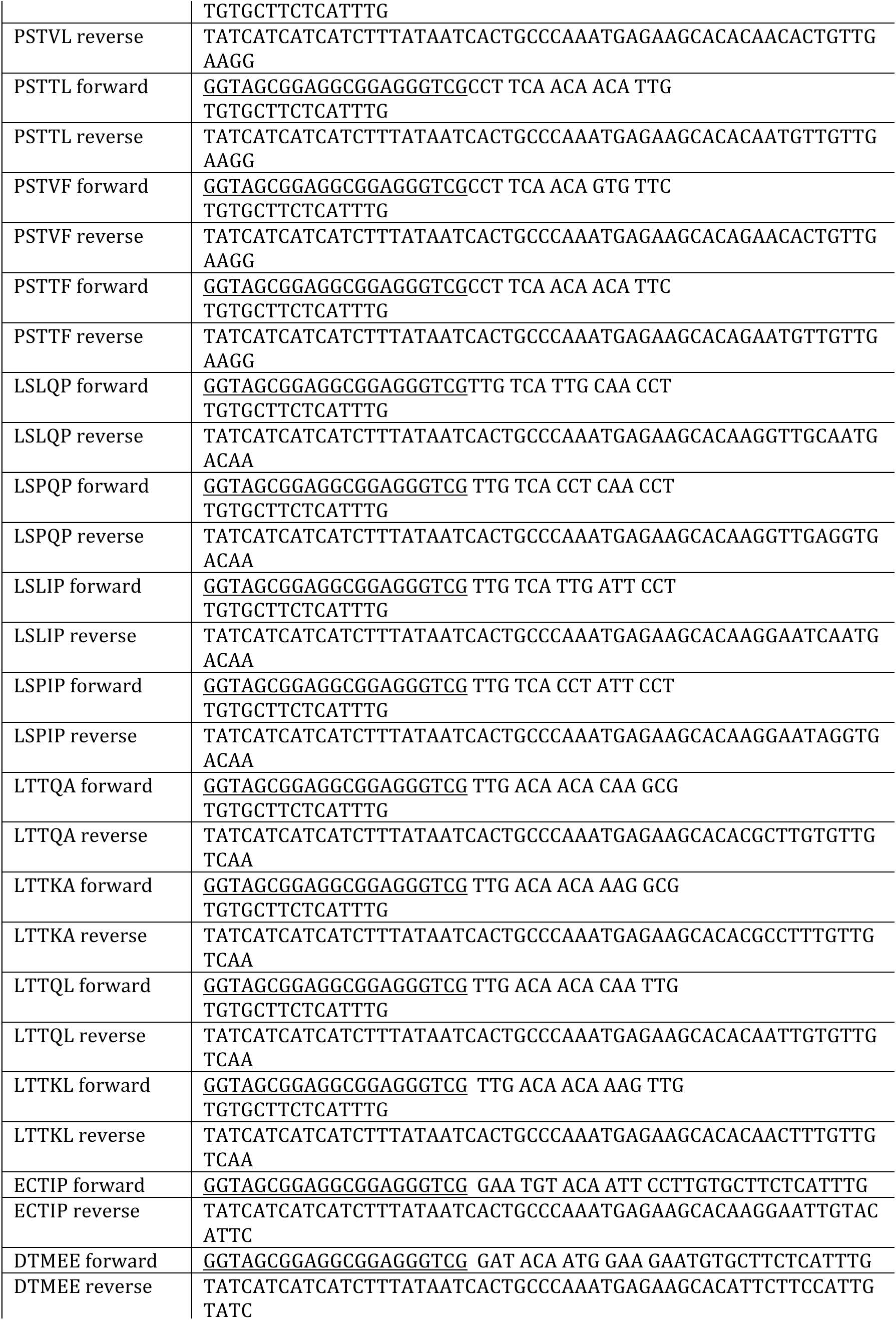
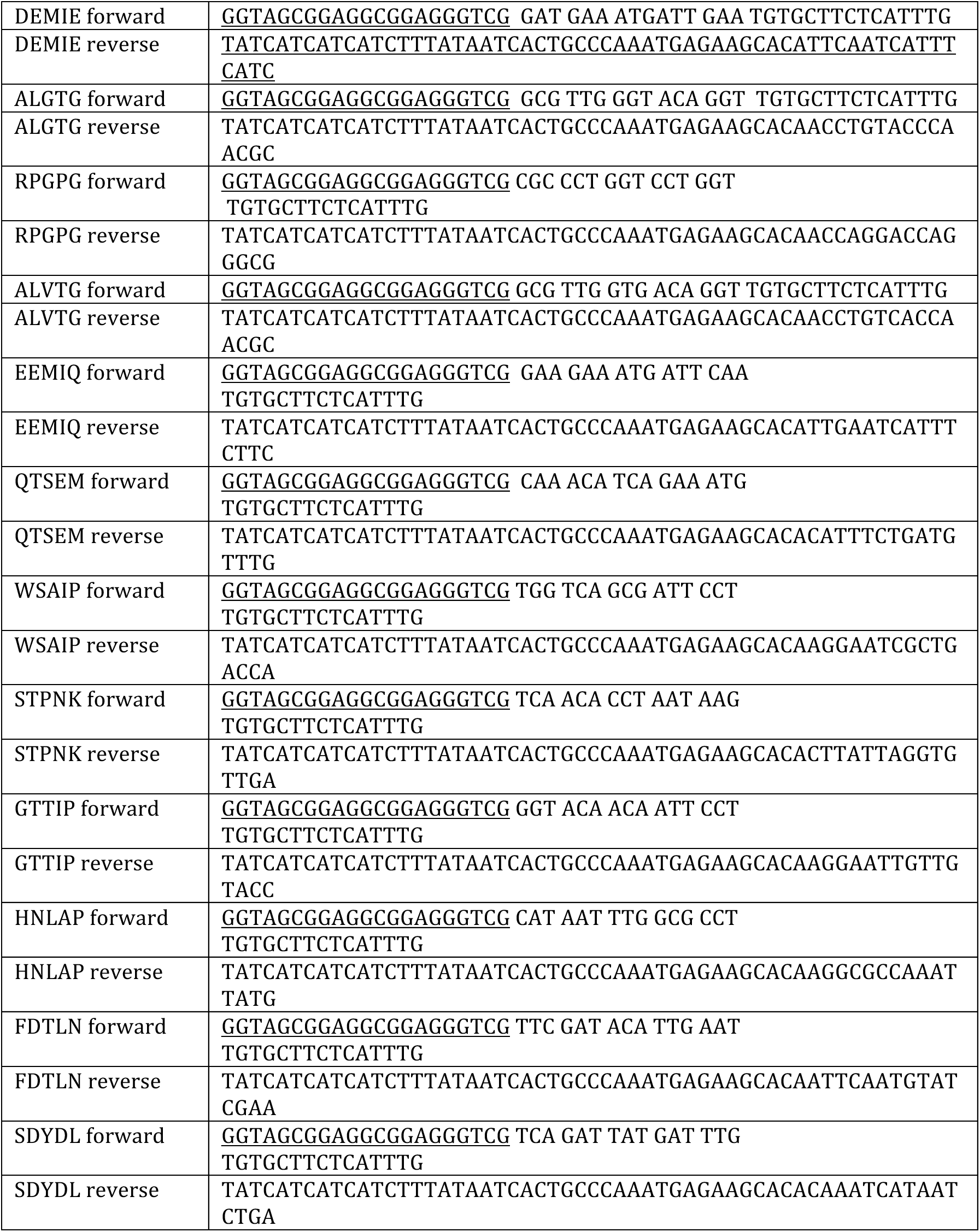
List of oligos for testing substrates in yeast surface display.

**7.**
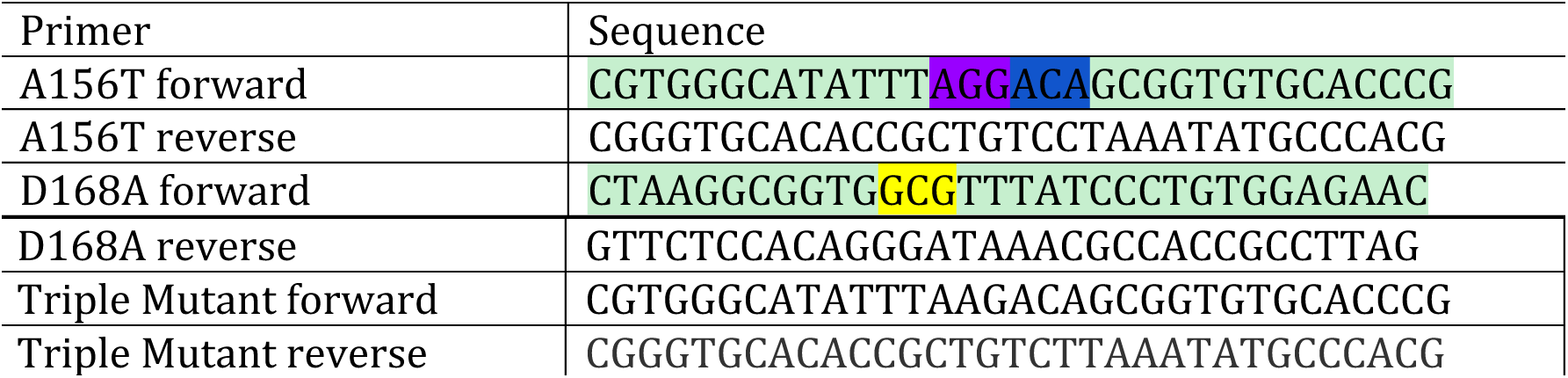
Primers to generate Drug Resistant Mutants.

**8.**
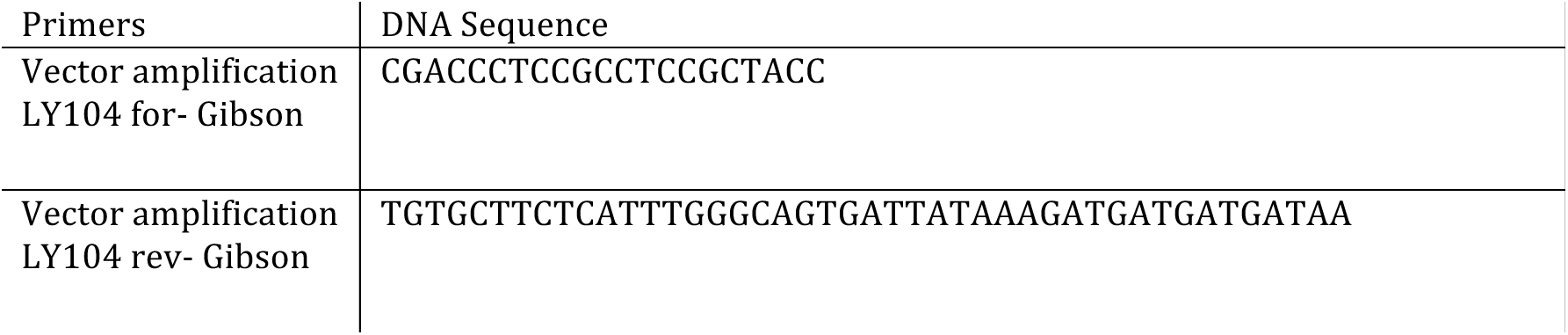
Vector amplification primers for YESS assay.

**9.** SVM parameter tuning: grid search for optimal boxconstraint and rbfsigma parameters. Average AUC is for each set of parameters run with an 80:20 split on the WT experimental full data set for 100 iterations. N/A is shown if the SVM did not converge with these parameters. A boxconstraint of 1 and rbfsigma of 10 was decided on for future calculations.

|  |  | boxconstraint |  |  |  |  |  |
| --- | --- | --- | --- | --- | --- | --- | --- |
|  | AUC | 0.01 | 0.1 | 1 | 10 | 100 | 1000 |
| rbfsigma | 0.01 | 0.5 | 0.5 | 0.5 | 0.5 | 0.5 | 0.5 |
|  | 0.1 | 0.5 | 0.5 | 0.5 | 0.5 | 0.5 | 0.5 |
|  | 1 | 0.8715 | 0.8718 | 0.872 | 0.872 | 0.8723 | 0.8721 |
|  | 10 | 0.9549 | 0.9811 | 0.9839 | 0.9829 | 0.9809 | 0.981 |
|  | 100 | 0.9695 | 0.9696 | 0.975 | 0.919 | 0.9825 | N/A |
|  | 1000 | 0.9691 | 0.9691 | 0.9693 | 0.9691 | 0.9748 | 0.9819 |

